# A mechanobiological computational framework of intestinal healing for sutureless surgical strategies

**DOI:** 10.64898/2026.09.24.753298

**Authors:** Syed Muhammad Zubair Shah Bukhari, Pietro Lenarda, René Thierry Djoumessi, Alessio Gizzi, Marco Paggi

## Abstract

Healing of the gastrointestinal (GI) tract following surgical interventions is governed by coupled mechanobiological processes and may lead to severe complications such as perforation or stricture formation. A first comprehensive mechanobiological modeling framework of GI healing is herein proposed. The framework addresses two clinically relevant applications of bioprinting: endoscopic treatment of GI tissue defects and bioprinting-assisted surgical anastomosis. Tissue remodeling is based upon a hyperelastic material model with fiber dispersion coupling the spatiotemporal reaction–diffusion dynamics of fibroblasts and cytokines with collagen deposition, reorientation, and plastic deformation. An active stress formulation drives fibroblasts and myofibroblast contraction while a multiplicative decomposition approach is adopted for collagen remodeling as a permanent change. The governing equations are discretised in time with an implicit Euler scheme, finite element in space and solved by a staggered algorithm implemented in the open-source platform FEniCS. Numerical simulations show that the initial collagen fibers content and orientation strongly influence fibroblast infiltration and scar contracture formation. The study quantifies the advantage of having a collagen bioprinted architecture aligned with tissue fibers, to reduce structural remodeling distortion. The model also shows that the recovery of mechanical integrity in GI anastomosis has a strong correlation with collagen deposition and luminal burst pressure.

## 1. Introduction

The gastrointestinal (GI) tract plays a central role in human health, and diseases affecting the digestive system represent a major clinical burden. Among them, colorectal cancer remains one of the leading causes of cancer-related mortality worldwide [1] with early detection and removal of precancerous lesions significantly improving patient survival rates [2]. Pathological conditions and surgical interventions can compromise the mechanical integrity of the intestinal wall, which can lead to severe complications such as perforation or leakage [3]. Intestinal wall healing typically requires weeks to months [4], and it is also subject to continuous exposure to luminal contents and biochemical agents. Therefore, understanding the mechanobiological mechanisms governing intestinal healing is essential for improving surgical outcomes and preventing postoperative failure.

Mechanobiological processes include a multitude of interactions between biochemical signaling, cellular activity, and tissue mechanics. Wound healing proceeds through four overlapping phases: hemostasis, inflammation, proliferation, and remodeling. During the later inflammatory phase, cytokines accumulate at the wound site by macrophages and attract fibroblasts, which proliferate and differentiate into myofibroblasts responsible for collagen synthesis during the proliferative phase. The newly deposited collagen is initially disorganized and progressively reoriented along the direction of maximum principal stretch. Fibroblasts and myofibroblasts generate contractile forces, while continued collagen deposition consolidates the contractile deformation, leading to irreversible contracture during the remodeling phase [5].

Diagnosis and treatment of GI lesions is routinely performed via endoscopy. Operative endoscopic procedures, such as mucosal resection, enable minimally invasive removal of mucosal neoplastic lesions [6]. The Endoscopic Submucosal Dissection (ESD) [7], in particular, includes removal of the submucosal layer by injecting saline to enable for dissection, as shown in Figure 1. These procedures improve patient outcomes due to their reduced invasiveness, still leading to post-interventional complications. As an example, endoscopic hemostatic techniques, such as styptic colloid spraying [8], can control bleeding but cannot restore the native three-dimensional (3D) structure of the GI wall, potentially compromising the healing process.

**Figure 1:**
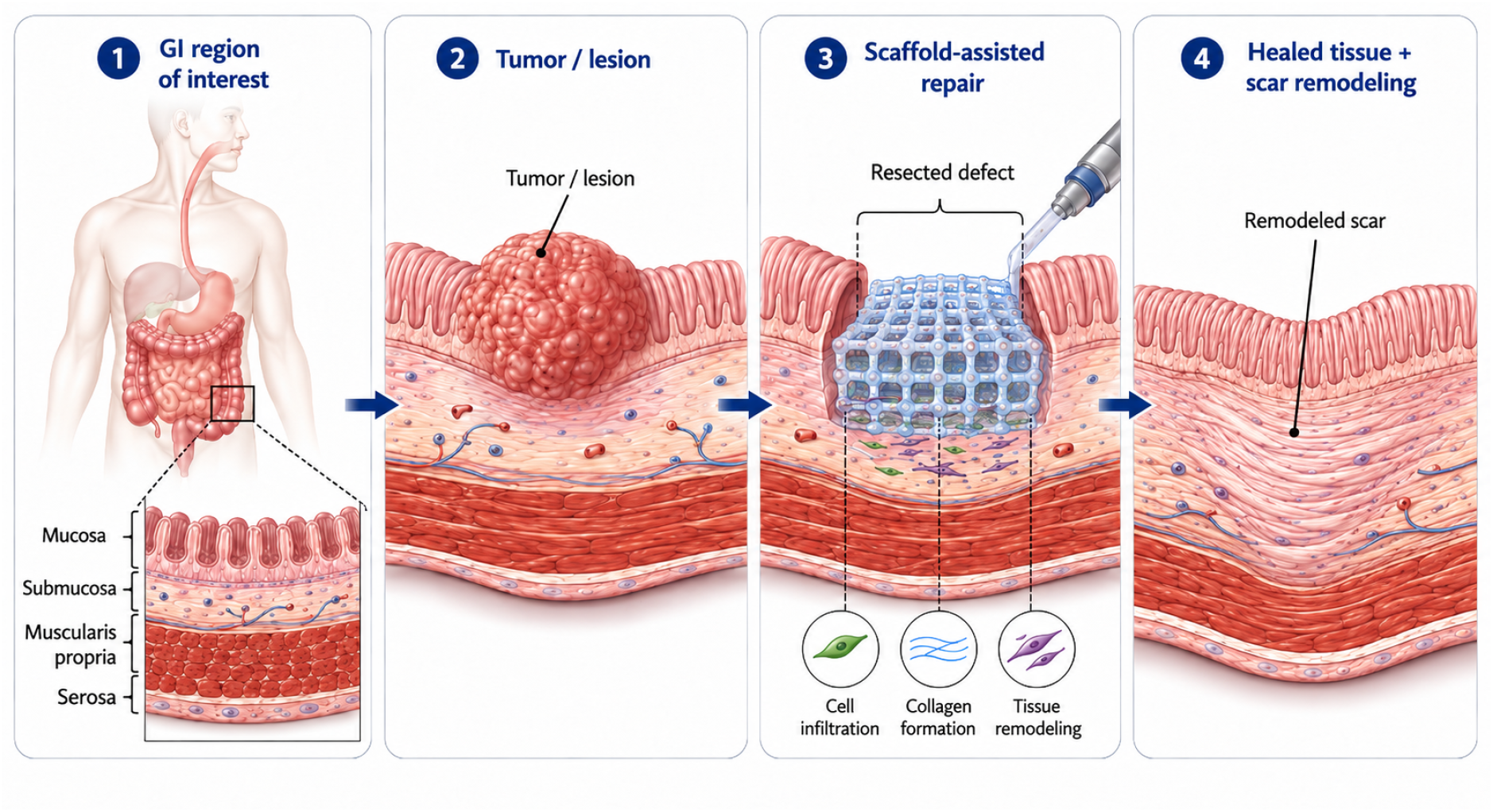
Schematic representation of the in situ GI tract intervention workflow. The illustration depicts the (1) region of interest with intestines, (2) lesion detection and dissection, (3) localized deposition of biomaterial via three-dimensional bioprinting within the wound region, and (4) subsequent healing and scar-remodelled tissue. This images has been generated with AI.

Postoperative GI wall healing may also lead to adverse mechanical outcomes. Excessive fibrotic remodeling can cause tissue stiffening and abdominal adhesions, where fibrous scar tissue abnormally connects adjacent organs or intestinal segments [9]. These conditions impair physiological intestinal motility and potentially induce to bowel obstruction. Clinical studies report that abdominal adhesions occur in more than 90% of patients following abdominal surgery [10]. To enhance tissue repair following endoscopic interventions, several regenerative strategies have been proposed, including tissue welding [11] and biomaterial-based therapies [12]. 3D-bioprinting has emerged as a promising approach for restoring damaged GI tissue by depositing biomaterials containing living cells and bioactive agents directly at the defect site [13]. However, the long-term mechanical stability of the scaffold (bio-printed implants) and their interaction with the host tissue during healing remain largely unexplored.

GI anastomosis is a surgical procedure performed in severe disease cases, involving the resection of a diseased intestinal segment followed by the reconnection of the remaining healthy tissue ends [14]. The postoperative healing of the anastomotic junction is a critical determinant of surgical success, as complications such as anastomotic leakage [15] or luminal stricture formation [16] can severely impair intestinal function and patient recovery. Clinical observations indicate that the postoperative days upto 7 represent the most vulnerable period for leakage [17], whereas long-term remodeling may lead to fibrotic contraction and luminal narrowing at the anastomotic site [18]. To evaluate the success of anastomotic surgery, the burst pressure test is commonly used to evaluate the mechanical strength [19]. To improve the mechanical strength of the anastomosis region and avoid leakage, bioadhesive materials are also used [20].

To the best of the authors’ knowledge, a comprehensive study investigating the coupled biochemical and mechanical effects of scaffold-based interventions in ESD, as well as anastomotic healing for the evaluation of burst pressure in the GI tract, has not yet been reported in the literature. Developing predictive computational models of intestinal healing is essential for capturing the coupled mechano-biological processes governing tissue repair. Compared to other biological systems, such as arteries [21], skin [22, 23], and breast [24], computational models of healing in the GI tract have received comparatively limited attention. In this work, we propose an active strain and active stress-based in silico mechanobiological framework for GI healing that captures the coupled interactions between biochemical signaling, cellular activity, and tissue mechanics for the ESD and anastomosis. The active strain accounts for contracture due to irreversible deformation, and active stress accounts for stress induced by fibroblasts and myofibroblasts. The model accounts for key processes, including infiltration of fibroblasts, cytokine concentration, collagen deposition, fiber reorientation, dispersion, and active contraction within a structurally constitutive description.

The main contributions of the present work are threefold. First, an established mechanobiological wound-healing formulation is adapted to intestinal wound healing by coupling it with a distributed four-fiber-family constitutive model of the intestinal wall, thereby accounting for its anisotropic mechanical behavior. Second, the resulting framework is applied to ESD to investigate the effects of biomaterial reinforcement and scaffold fiber orientation on the spatiotemporal healing response and wound contracture. Third, intestinal anastomotic healing is investigated by assessing the progressive recovery of mechanical integrity under luminal pressurization, using a stress-based criterion as a computational surrogate for burst resistance.

The manuscript is organized as follows: section 2 presents the mechanobiological healing model of the intestinal wall; section 3 describes the numerical strategies and their implementation; section 4 investigates the applications of the in-silico model and section 5 conclude the work with limitations and future directions.

## 2. Mechanobiological constitutive modeling of intestinal wall

This section presents the governing equations for the coupled mechanobiological system, integrating biochemical transport and mechanical deformation with microstructural remodeling through both active stress and active strain formulations. We employ standard mathematical notation, where scalars are denoted by (*a*), vectors by (***a***), and second-order tensors by (***A***), with (***A***^*T*^) indicating the tensor transpose. Consistent with continuum mechanics conventions, the scalar product between vectors is denoted by (·), the double contraction of tensors by (:), and the dyadic product by (⊗). Differential operators are defined as follows: ∇ represents the gradient, ∇· the divergence, and ∇^2^ the Laplace operator. This notation provides a consistent mathematical framework for describing the multiphysics coupling between biochemical fields, finite-strain mechanics, and tissue remodeling during colonic wound healing.

### 2.1. Finite kinematics

The kinematics of deformable GI tissue are formulated within the classical framework of continuum mechanics under finite elasticity [25]. Let ***X*** and ***x*** denote the material coordinates in the reference (Ω_0_) and current (Ω_*t*_⊂ ℝ^*d*^, *d* = 3) configurations, respectively. The deformation mapping ***ψ*** : Ω_0_ → Ω_*t*_ defines the deformation gradient ***F*** = *∂****x****/∂****X*** with Jacobian *J* = det ***F*** *>* 0. The right and left Cauchy–Green deformation tensors are given by ***C*** = ***F***^*T*^ ***F*** and ***B*** = ***F F***^*T*^, respectively.

In the following, we identify with *ρ*(***X***, *t*) fibroblast density, *c*(***X***, *t*) cytokine concentration, and *ϕ*(***X***, *t*) collagen density; collagen matrix is further defined through the fiber dispersion and preferred fiber orientation *κ*(***X***, *t*) and ***n***_*i*_(***x***, *t*), respectively, where the index *i* represents a set of fiber directions.

To represent the multiphysics coupling between tissue mechanics and cellular activity, a multiplicative decomposition of the deformation gradient is adopted, allowing for a separation of elastic deformation from plastic deformation under finite strains:

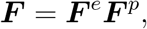

where, ***F***^*e*^ and ***F***^*p*^ denote the elastic and plastic components, respectively. The plastic deformation tensor ***F***^*p*^ thus encodes local irreversible remodeling induced by fibroblast and myofibroblast contractility, defined through scalar stretch fields 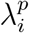 and preferred fiber orientations. This anisotropic plastic deformation tensor that accounts for permanent deformation [26, 27] can be expressed as:

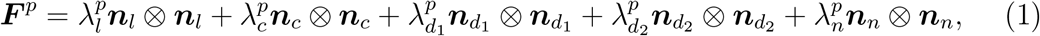

with ***n***_*l*_ and ***n***_*c*_ the longitudinal and circumferential directions, respectively, with ***n***_*d*1_ and ***n***_*d*2_ the diagonal fiber directions, and ***n***_*n*_ represents the normal unit vector orthogonal to the ***n***_*l*_–***n***_*c*_ plane. Circumferential and longitudinal fiber families are representative of smooth muscle cells layers which are present in the GI tract (stratum longitudinale and stratum circulare), while diagonal fibers correspond to collagen fibers reinforcement [28, 29, 30], in the submucosal layer. These directions are in the reference configurations.

### 2.2. Biochemical model

The biochemical model describes the spatiotemporal evolution of fibroblast density and cytokine concentration, via a coupled reaction-diffusion system (*ρ*(***X***, *t*) = *ρ, c*(***X***, *t*) = *c*). These fields capture the key biochemical interactions driving tissue deformation and remodeling during healing [5] as:

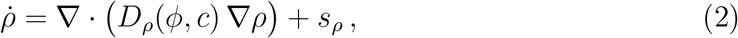

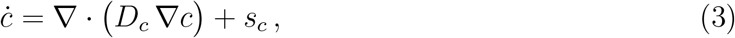

The cytokine diffusivity *D*_*c*_ is assumed constant, whereas the fibroblast diffusivity *D*_*ρ*_(*ϕ, c*), depends on *c* and nonlinearly of local collagen content *ϕ* and [24] and is expressed as:

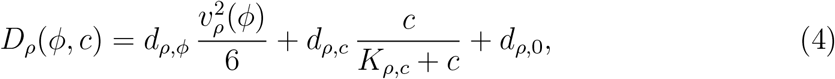

with *d*_*ρ,ϕ*_, *d*_*ρ,c*_, and *d*_*ρ*,0_ the collagen-dependent, cytokine-dependent, and baseline diffusion coefficients, respectively, considering a Michaelis–Menten-type saturation kinetics where *v*_*ρ*_(*ϕ*) represents the fibroblast migration speed as a function of collagen density, estimated from in-vivo wound-healing data [31, 32, 33].

The reaction terms account for production and degradation of fibroblasts and cytokines:

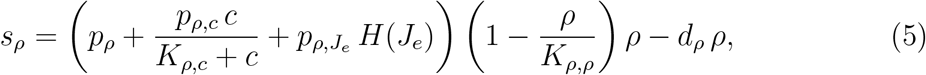

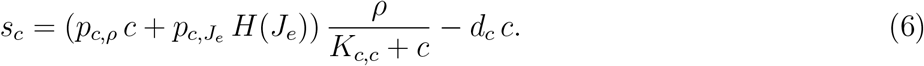

The mechanical feedback loop via the sigmoidal function *H*(*J*_*e*_), linking the elastic volumetric change, *J*_*e*_ = det(***F***^*e*^), with biochemical activity.

### 2.3 Constitutive mechanical model

The GI wall is approximated as a compressible, anisotropic hyperelastic material whose passive mechanical response depends on the elastic component of deformation. Under the multiplicative decomposition ***F*** = ***F***^*e*^***F***^*p*^, the elastic Jacobian is *J*_*e*_ = det(***F***^*e*^), and the compressibility of the elastic response is enforced through the volumetric energy. The strain–energy density is additively split into isotropic (matrix), volumetric, and anisotropic parts, collectively scaled by the local collagen content *ϕ*:

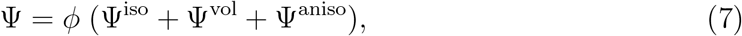

where *ϕ* is the collagen content. This scaling provides a phenomenological coupling between collagen density and mechanical recovery during healing. Collagen content is treated as a proxy for overall tissue mechanical integrity.

The isotropic matrix response follows a Neo-Hookean form:

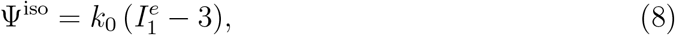

where *k*_0_ is the stiffness of the ground matrix, and the first invariant is defined as 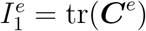 where ***C***^*e*^ = ***F***^*eT*^ ***F***^*e*^.

The volumetric part is used in the strain energy function for a compressible material

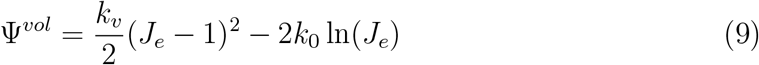

where *k*_*v*_ is the bulk modulus controlling the resistance to elastic volume changes. Large values of *k*_*v*_ increasingly penalize volumetric deformation, recovering the nearly incompressible response.

Four fiber families are considered: longitudinal (*l*), circumferential (*c*), and two diagonal fiber families (*d*_1_, *d*_2_). Collagen-fiber dispersion is introduced in the strain energy function through the dispersion parameter *κ* [34] as:

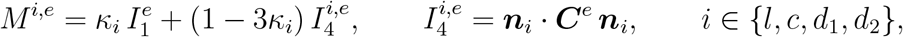

where 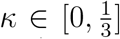 denotes the dispersion parameter (*κ* = 0 for perfectly aligned fibers and 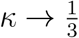representing the highest admissible statistical dispersion, and the vectors ***n***_*i*_ denote the preferred fiber directions. The passive anisotropic contribution is expressed as

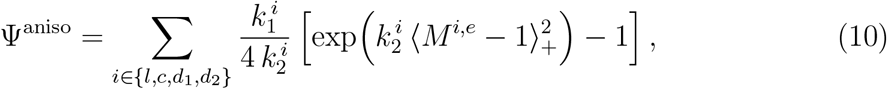

where 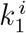 (stress-like) and 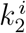 (dimensionless) are material constants controlling fiber stiffness and nonlinearity, respectively. The Macaulay bracket ⟨*y*⟩_+_ = max(*y*, 0) enforces the tension–compression switch, suppressing fiber resistance under compression.

According to the given prescriptions, the first Piola–Kirchhoff passive stress tensor is derived as:

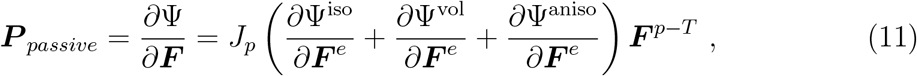

where *J*_*p*_ = det(***F***^*p*^). The total stress will be completed in the next section with the introduction of the active stress acting in the wound region due to the traction of the fibroblasts and myofibroblasts.

### 2.4. Mechanobiological coupling

The mechanobiological coupling describes how biochemical activity (cell proliferation and cytokine signaling) interacts with mechanical deformation through two principal mechanisms: (i) a mechanosensing logistic function that activates beyond homeostatic deformation, and (ii) an active stress contribution generated by fibroblast and myofibroblast contractility along preferred fiber directions. These terms provide the bidirectional coupling between the biochemical and mechanical fields, closing the feedback loop that governs tissue remodeling.

Mechanical activation is introduced through a smooth logistic function that depends on the elastic Jacobian *J*_*e*_ = det(***F***^*e*^). The function captures the transition between physiological (homeostatic) and mechanically activated states:

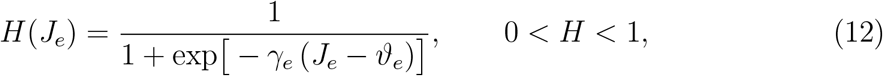

where *γ*_*e*_ controls the shape of mechanosensing curve and *ϑ*_*e*_ is the deformation threshold for mechanosensing activation, both values for these parameters are reported in Table E.3. This functional form ensures a gradual activation of the cellular response once the deformation departs from the homeostatic configuration.

The active stress represents the contractile contribution generated by fibroblast and myofibroblast activity, modulated by local biochemical and mechanical cues. A structural tensor approach is adopted to capture fiber-aligned contraction over multiple fiber families. The active stress is expressed as:

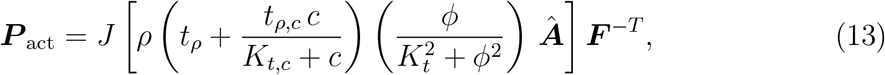

where, *t*_*ρ*_ and *t*_*ρ,c*_ govern the baseline contractility and enhanced cytokine, respectively, while *K*_*t,c*_ and *K*_*t*_ are saturation constants controlling the Michaelis–Menten-type response.

The directionality of active contraction is defined through a generalized structural tensor that aggregates the contributions of the longitudinal (*l*), circumferential (*c*), diagonals (*d*_1_, *d*_2_), and normal (*n*) directions:

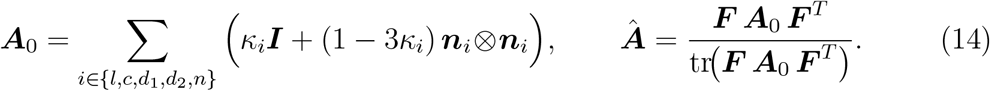

The resulting *Â* provides a normalized structural measure of anisotropy that defines the preferential directions for active contraction in the current configuration.

The coupling is done in two way for the forward coupling with the biochemical and microstructure via ***P*** _*act*_, while the feedback coupling is done by the mechanosensing activation function *H*(*J*_*e*_).

### 2.5. Micro-structural fields

The micro-structural fields are coupled to the mechanics and biochemical fields and locally evolve through a set of ordinary differential equations (ODEs). We consider collagen density *ϕ*, fiber dispersion parameter *κ*, fiber orientation ***n***_*i*_, and plastic stretches 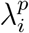 as local fields. The local remodeling parameters used are as shown in Table E.3.

#### 2.5.1. Collagen density

The fibroblasts are the main contributors to the deposition of collagen in the wound area. The first term in right side have base collagen production, production enhanced by cytokines and then by mechanotransduction. The second term is the collagen degradation, having natural decay and decay influenced by fibroblasts and cytokines:

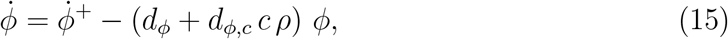

where:

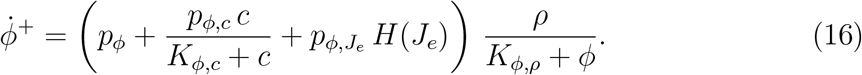

### 2.5.2. Fiber dispersion (κ)

We assume all collagen fiber families share the same dispersion level 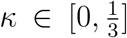, which is a simplification and limitation of the model. Its evolution is driven by collagen accumulation and elastic principal stretches:

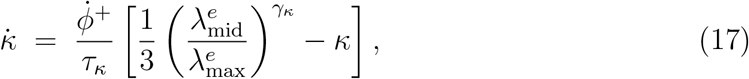

where 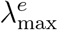 and 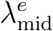 are respectively the maximum and second largest eigenvalue of the tensor ***C***^*e*^, *γ*_*κ*_ *>* 0 controls the sensitivity to the spread of principal stretches. The fiber dispersion is ruled by the ratio between the in-plane principal stretches, as the ratio 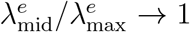, meaning isotropic deformation, then *κ* → 1*/*3 while if 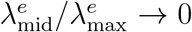 then *κ* → 0.This approach is similar to the approach used in [35, 36, 37], where the driving force is also the ratio of the principal stretches.

#### 2.5.3. Fiber reorientation

Fibers remodeling is assumed to be ruled by the direction of maximum principal elastic stretch. A projection-based evolution law keeping unit length is based on the works by [38, 39] as:

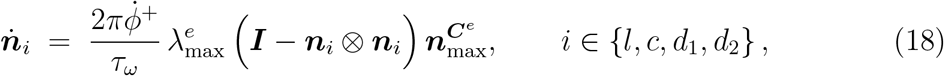

where 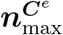 is the eigenvector of ***C***^*e*^ associated to the largest eigenvalue 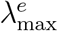 and *τ*_*ω*_ is the characteristic time. These evolving microstructural variables directly update fiber stretches, anisotropic invariants, and active stress generation introduced previously (see subsection.2.4).

#### 2.5.4. Plastic stretch

For each direction the plastic stretch is evolving as:

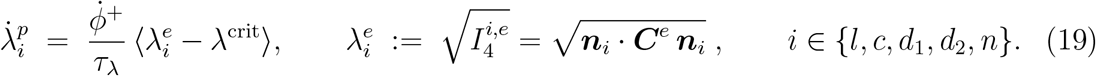

The plastic stretch accumulates only when collagen is deposited 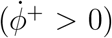 and the elastic stretch 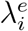 leaves the physiological deadband [0.995, 1.005]. Within this range, it is assumed that no plastic evolution occurs [5]. The resulting plastic stretches 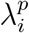 contribute to the plastic tensor ***F***^*p*^ introduced earlier for permanent deformation.

## 3. Numerical Implementation

### 3.1. Strong form

We consider the reference configuration Ω_0_ ⊂ R^*d*^ (*d* = 3) and the time domain [0, *T*]. The quasi-static mechanical equilibrium is formulated with displacement. The unknown field is the displacement ***u*** : Ω_0_ × [0, *T*] → ℝ^*d*^. The strong form of the governing equation reads as follows:

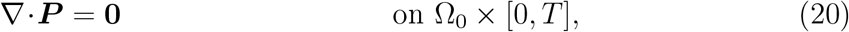

The boundary *∂*Ω_0_ is divided into displacement and traction parts, Γ_*D*_ and Γ_*N*_, such that Γ_*D*_ ⋂ Γ_*N*_ = 0 and Γ_*D*_ ⋃ Γ_*N*_ = *∂*Ω_0_.

Boundary conditions are prescribed as:

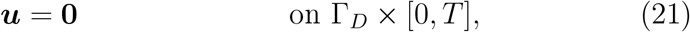

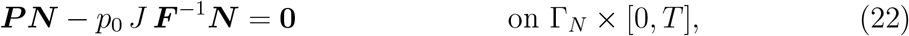

where *N* is the outward unit normal in the reference configuration. The displacement Dirichlet boundary conditions are imposed on Γ_*D*_ as in Eq. (21), where the nodes are fixed and no motion is permitted. These constraints prevent rigid body translation and rotation of the tissue domain. On the remaining portion of the boundary, Γ_*N*_ as in Eq. (21), a uniform pressure load is applied to represent the internal material contents (digesta) within the lumen.

For the biochemical problem we impose homogeneous Neumann (no-flux) conditions on the lumen–wall boundary. This enforces zero net transport of cells and cytokines across the boundary in the current configuration:

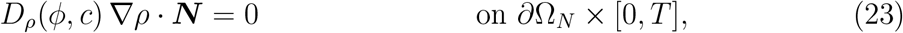

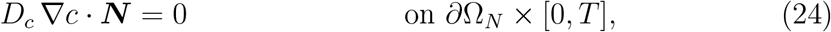

where *∂*Ω_*N*_ is the Neumann boundary. Equations (23)–(24) state that neither fibroblasts nor cytokines cross *∂*Ω_*N*_.

### 3.2. Weak variational form

#### 3.2.1 Mechanical Weak form

The displacement is defined in the function spaces:

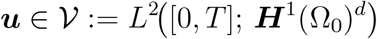

Virtual displacements and test functions are defined on the spaces vanishing on the Dirichlet boundary:

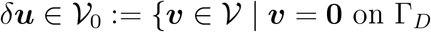

The weak problem reads: Find ***u*** ∈ V such that, for all *δ****u*** ∈ V_0_:

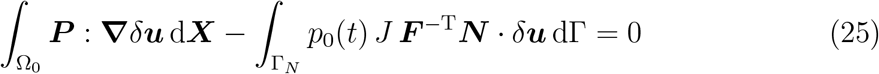

Here ***P*** is the total first Piola–Kirchhoff stress tensor, ***N*** is the outward unit normal vector on Γ_*N*_ ⊂ *∂*Ω_0_, *p*_0_(*t*) denotes the prescribed luminal pressure, *J* = det(***F***) is the deformation gradient.

#### 3.2.2. Biochemical weak form

Let *ρ*(***X***, *t*) and *c*(***X***, *t*) be the unknowns of the strong form in equation (2) and (3) with initial conditions *ρ*(***X***, 0) = *ρ*_0_(***X***), *c*(***X***, 0) = *c*_0_(***X***), and boundary conditions (e.g., no–flux) as in equations (23)–(24).

The biochemical fields are defined in the Sobolev space:

(*ρ, c*) ∈ W^2^ := [*L*^2^([0, *T*]; *H*^1^(Ω_0_))]^2^. Let the associated test functions vanish on Dirichlet boundaries: 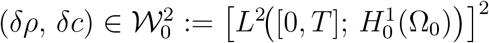.

Multiply the strong form (2) and (3) by the test functions (*δρ, δc*), integrate over Ω_0_, using the divergence theorem and the no–flux condition, the variational problem reads: Find (*ρ, c*) ∈ W^2^ such that:

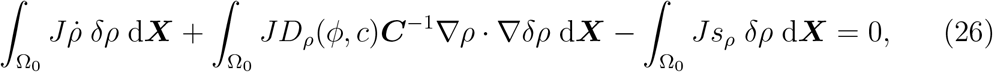

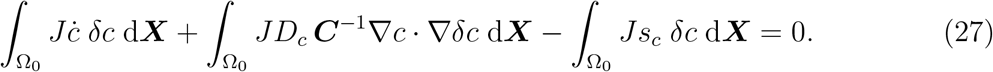

#### 3.2.3. Micro-structural field weak form

Multiply the strong form of each remodeling equation by its corresponding test function and integrate over Ω_0_. The weak problem reads: Find the vector of unknowns defined in a suitable weak vector space:

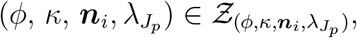

such that, for all test functions defined in the corresponding vector space:

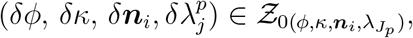

where indices *i* and *j* run respectively in the sets of fiber families defined as: *i* ∈ {*l, c, d*_1_, *d*_2_} and *j* ∈ {*l, c, d*_1_, *d*_2_, *n*}.

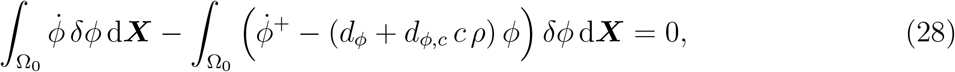

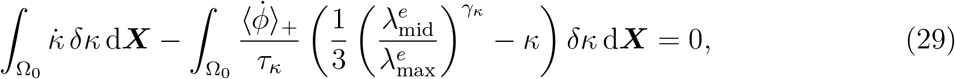

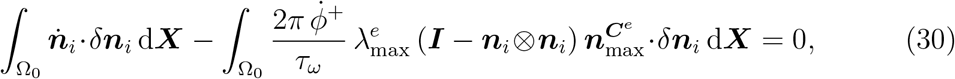

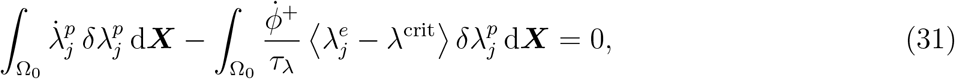

### 3.3. Finite element discretization

The computational domain was discretized using unstructured tetrahedral finite elements. All governing equations were approximated using Lagrangian shape functions with a ℙ_2_–ℙ_1_ formulation, where quadratic interpolation (ℙ _2_) was used for displacement, and linear interpolation (ℙ_1_) for fibroblasts and other fields. Simulations were performed in the open-source FEniCS framework [40], which provides automatic differentiation for consistent Jacobian evaluation.

A staggered solution scheme was employed to sequentially solve the mechanical, biochemical, and remodeling problems. Each nonlinear system was solved using the Newton–Raphson method, and the resulting linearized equations were resolved with the direct `MUMPS` solver. This approach allows robust coupling between the mechanical response, biochemical transport, and microstructural remodeling processes in the mechanobiological healing model.

## 4. Numerical experiments

Prior to the healing simulations, the fiber reorientation was validated as described in Appendix Appendix B (Figure B.15). We then performed intestinal healing simulations for endoscopic cases and the anastomosis case.

### 4.1. Case study I: Endoscopic submucosal Dissection of the intestinal tract

In the endoscopic submucosal dissection (ESD) case, an idealized cylindrical geometry is adopted, with a length of 100 mm, an inner diameter of 22 mm, and an outer diameter of 24 mm as shown in Figure 2 (a). The clinical ESD studies report lesion diameters ranging from a few millimeters to several centimeters, with early-stage neoplasms frequently measuring around 10 mm. Lesions smaller than 10 mm are commonly classified as small, whereas lesions between 10 and 20 mm are considered medium-sized and are routinely treated using ESD, particularly when en-bloc resection is required [41, 42]. Accordingly, this work selected a 10 mm wound diameter representative ESD defect. This choice reflects common clinical practice for localized intestinal lesions confined to the submucosal layer and is appropriate for the scope of the present study [43].

**Figure 2:**
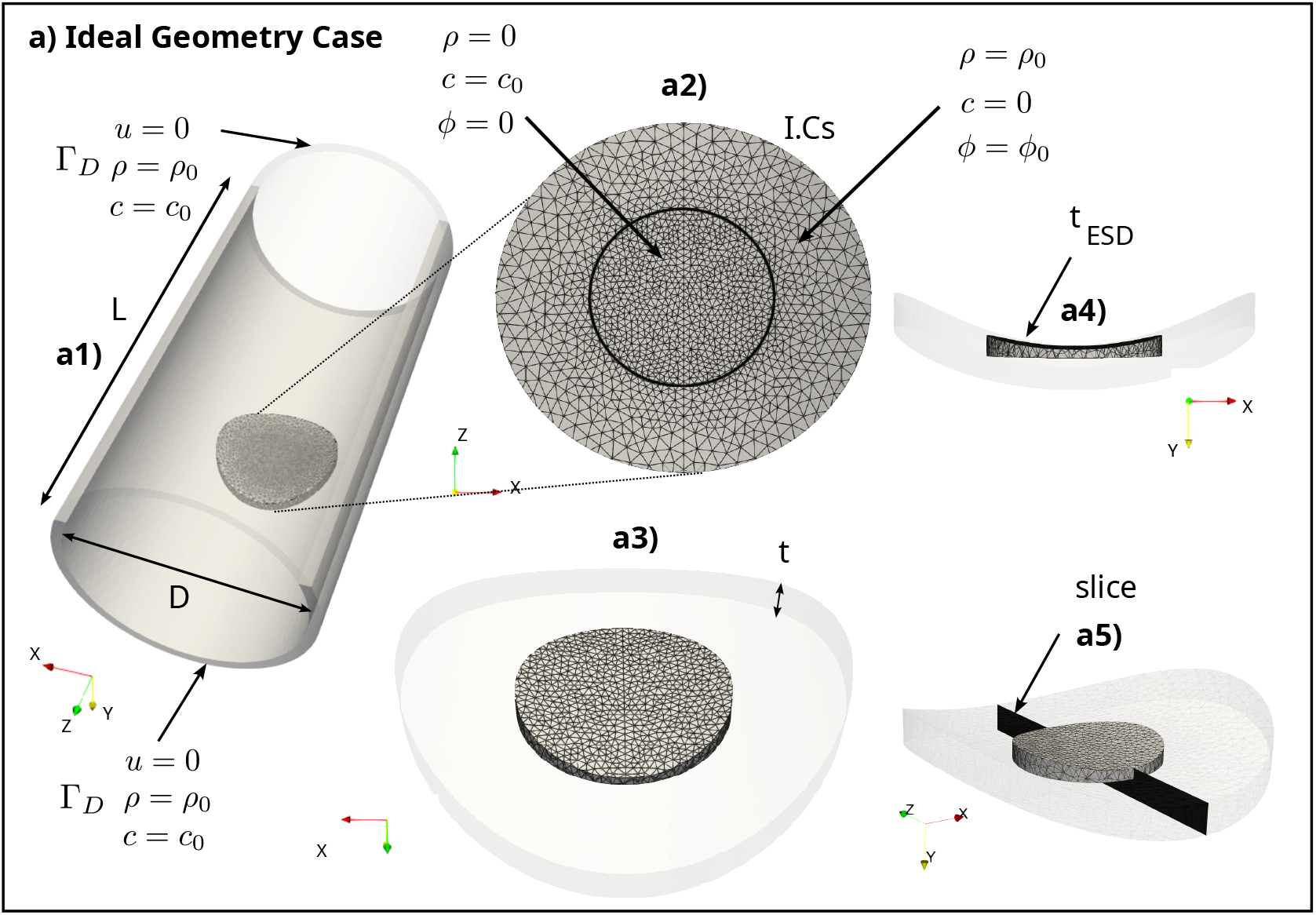
Schematic representation of the (a) idealized geometry (a1) The cylindrical geometry with length *L* and diameter *D*, where the wound and surrounding tissue patch are shown with opacity of whole geometry; Dirichlet boundary conditions are applied at the cylinder edges. (a2) Application of the initial conditions. (a3) Definition of the wound region. (a4) Specification of the wound thickness. (a5) Cross-sectional slice of the wound and surrounding tissue illustrating the local thickness reduction.

The wound region is located at the center of the inner surface of the cylindrical domain and is characterized by a diameter of 10 mm and a thickness of 1 mm. The intestinal wall is modeled using a layered fiber architecture comprising four families of fibers embedded at each material point. Among these, the diagonal submucosal collagen fibers ***n***_*d*1_, ***n***_*d*2_ are considered more biologically active during the healing process following ESD due to major collagen content in this layer, whereas the longitudinal and circumferential smooth muscle fibers ***n***_*l*_, ***n***_*c*_ are assumed inactive in terms of active contraction during the healing process because they are not subject to resection. Nevertheless, all fiber families contribute mechanically to load bearing and stress redistribution within the intestinal wall.

The boundary conditions are applied at both ends of the cylindrical domain, prescribing zero displacement, a maximum fibroblast density, and zero cytokine concentration as shown in Figure 2 (a1). Initially, the wound region is characterized by zero fibroblast and collagen densities, while the cytokine concentration is set to its high value to demonstrate high inflammation. Outside the wound region, fibroblast and collagen densities are initialized at their maximum value to show the early proliferation stage, where chemical attractors are inside the wound, with the cytokine concentration set to zero as shown in Figure 2 (a2).

Based on this framework, two case studies are investigated for the ESD scenario: the effect of the initial collagen density of the 3D-printed implant and the influence of its initial fiber orientation on healing and mechanical remodeling after surgery. The schematics of the no-fill case (a) and the scaffold-filled cases with fiber orientations (b) and (c) can be found in Appendix F. The initial fiber is generated using a modified rule-based method explained in our previous works [30]. The mechanobiological parameters used in our work can be found in Appendix E.

#### 4.1.1. The role of initial collagen density of the 3D bio-printed implant

The mechanobiological field variables are extracted at the center of the wound for the no-fill configuration and for scaffold-assisted cases with varying initial collagen content, as shown in Figure 3 (a)-(d). The fibroblast density *ρ* represents the primary cellular driver of collagen production and active stress generation within the model.

**Figure 3:**
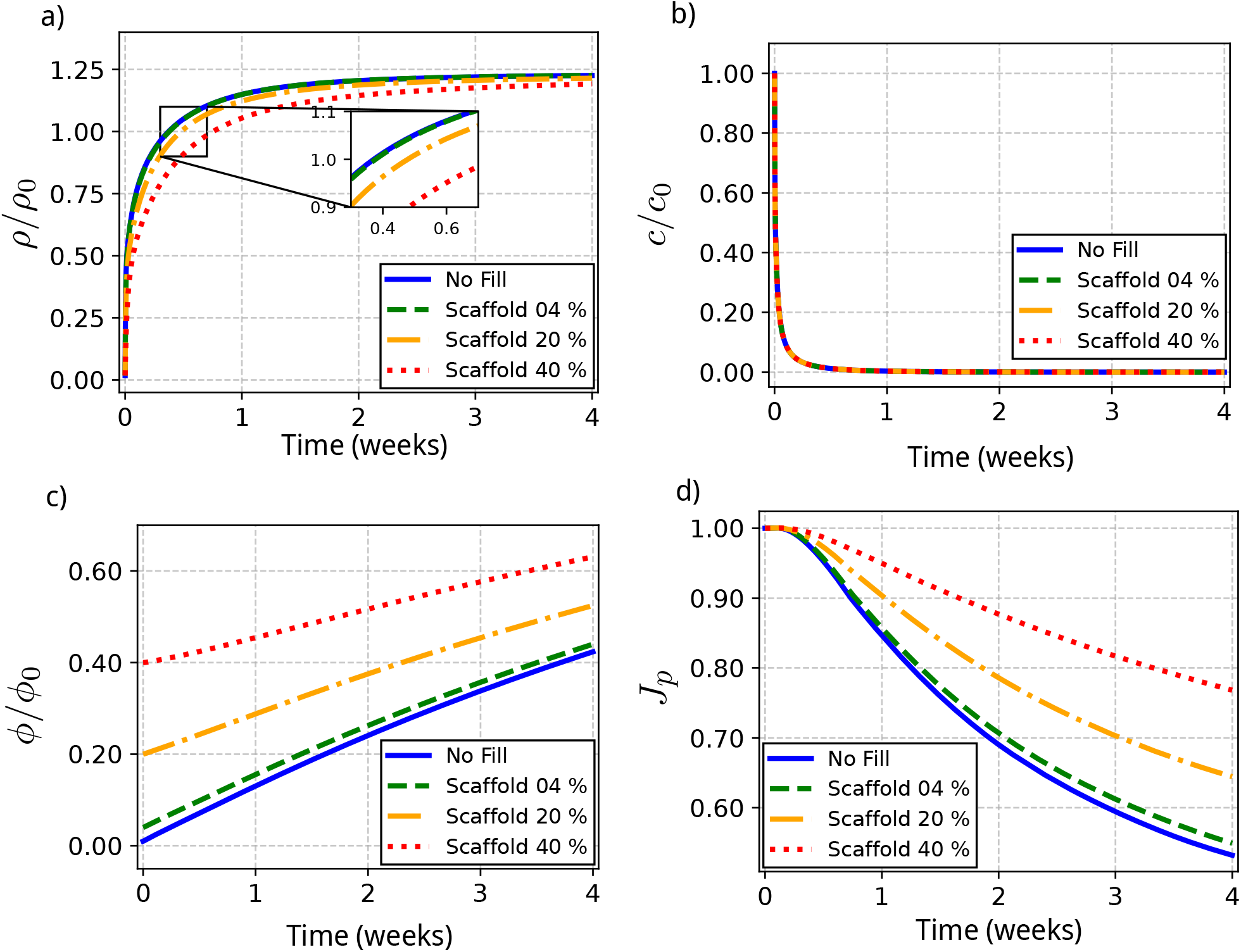
The evolution of (a) fibroblast infiltration, (b) cytokine concentration, (c) collagen density, and (d) contracture at the center of the wound for endoscopic submucosal resection cases with no fill and with collagen scaffolds of different densities, over four weeks.

Fibroblast infiltration is higher in the no-fill and 4% scaffold case, whereas the infiltration decreases progressively as the initial collagen content increases from 20% to 40%. The fibroblast density in the no-fill and scaffold 4% cases is nearly identical during the early phase and reaches the physiological tissue fibroblast density at approximately 0.38 weeks. In the subsequent weeks, the density stabilizes and attains a slightly higher value than the physiological baseline.

In contrast, the 20% and 40% scaffold configurations reach the same physiological level of healthy tissue at approximately 0.47 and 0.75 weeks, respectively. During the initial 1.5 weeks, a clear separation is observed between all configurations; however, beyond the third healing period, the fibroblast densities become very close to each other. This behavior is attributed to the initial collagen content within the wound region. A denser collagen network mechanically resists fibroblast infiltration, an effect that is explicitly captured in the model. Cytokine concentration decreases rapidly during the first three days and converges to an asymptote at the end of the first week, as shown in Figure 3 (b). As cytokines act as chemo-attractants for fibroblasts, their reduction limits fibroblast infiltration after one week. Collagen deposition initially exhibits an approximately linear trend for all cases, with the no-fill and 4 % scaffold cases showing nearly identical close behavior as shown in Figure 3 (c) due to initial close content. These results indicate that fibroblast infiltration is primarily governed by cytokine signaling and the initial collagen content. Figure 3 (d) shows that the contracture at the center of the wound vary from 0.52, 0.54, 0.65, and 0.77 for no fill, 4%, 20% and 40% collagen content. This initial collagen content supports the matrix and redistributes the stress generated by the myofibroblast to avoid the wound gimpling inward.

After analysing the temporal evolution of the mechanobiological variables at the wound center, the full three-dimensional spatio-temporal distributions are examined to elucidate how fiber re-alignment governs tissue regeneration and structural remodeling. Figure 4 and Figure 5 present the distributions of fibroblast density (*ρ*), cytokine concentration (*c*), collagen density (*ϕ*), and contracture (*J*_*p*_) for the no-fill and 40% collagen scaffold configurations at week 0 and 4 respectively.

**Figure 4:**
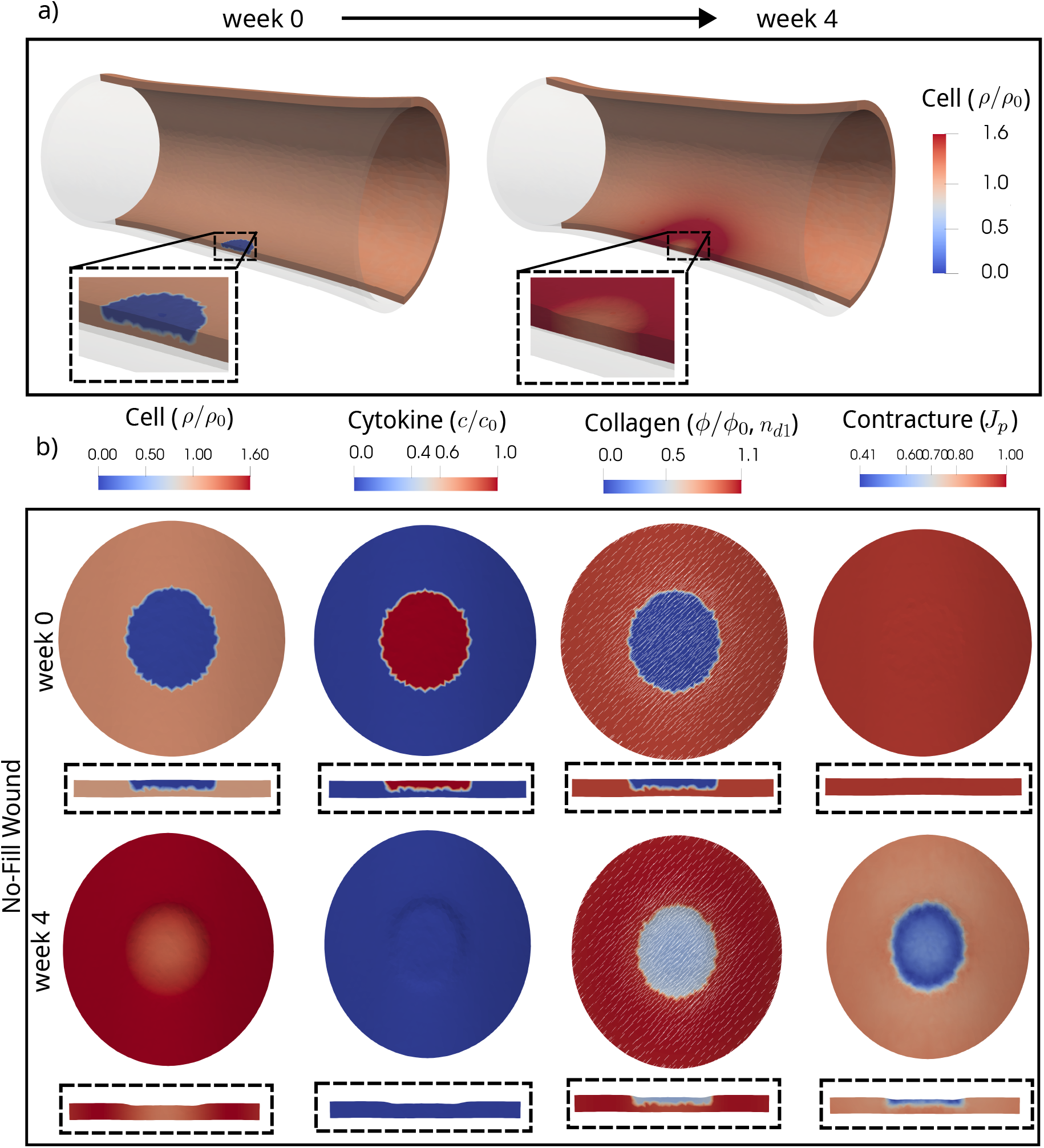
(a) Temporal solution illustrating changes in the wound region and cell production and infiltration in the wound region. (b) Extracted 3D subsection of the geometry showing the temporal evolution of fibroblast density, cytokine concentration, collagen density, and wound contracture for the no-fill collagen case shown at weeks 0 and 4.

**Figure 5:**
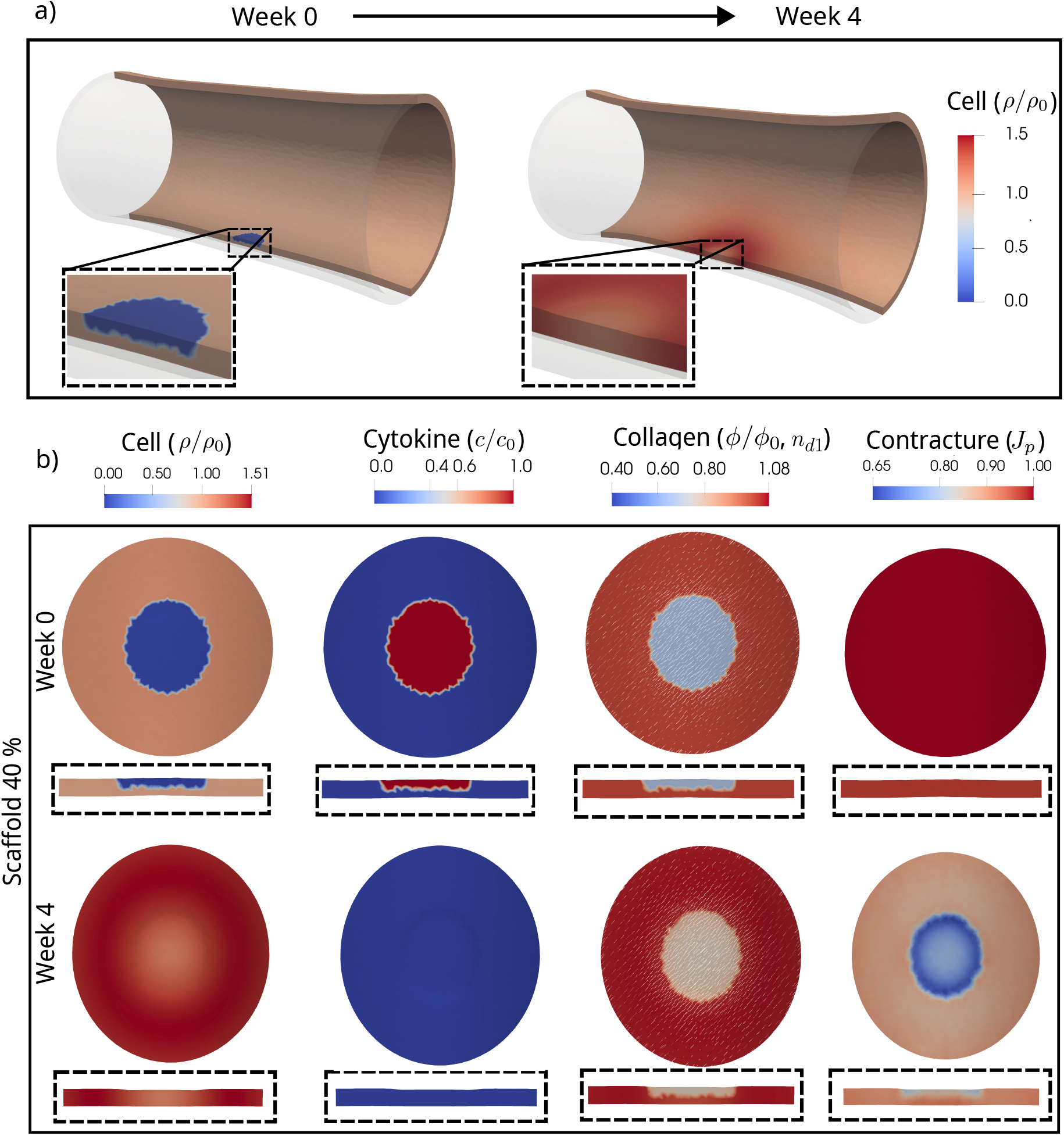
(a) Temporal solution illustrating changes in the wound region and cell production and infiltration in the wound region. (b) Extracted 3D subsection of the geometry showing the temporal evolution of fibroblast density, cytokine concentration, collagen density, and wound contracture for the 40 % scaffold aligned with the physiological angle, shown at weeks 0 and 4.

As shown in Figure 4 (a), zoomed wound region show contracture effect on the whole geometry at week 4. The clipped 3D geometry highlights the spatial gradients across the wound–healthy interface as shown in Figure 4 (b). By week 4, the wound and surrounding tissue show an elevated normalized fibroblast density of approximately *ρ/ρ*_0_ = 1.6, compared with the physiological value *ρ/ρ*_0_ = 1.0, with more infiltration due to less collagen and enhanced proliferation with cytokines and mechanotransduction. As fibroblasts deposit extracellular matrix within the wound region, the collagen density increases from *ϕ* = 0.01 at day 1 to approximately *ϕ* = 0.4 by week 4. Significant permanent contraction *J*_*p*_ is also observed in the wound and surrounding tissue. Here, *J*_*p*_ = 1 denotes no permanent volume change, *J*_*p*_ *<* 1.0 denotes contracture, and *J*_*p*_ *>* 1.0 denotes local stretch. Initially, *J*_*p*_ = 1 throughout the whole geometry, whereas by week 4, *J*_*p*_ = 0.41, indicating substantial permanent contracture corresponding to approximately 59% local volume reduction relative to the initial configuration.

In contrast, the 40% collagen scaffold configuration substantially attenuates the structural distortion, as shown in Figure 5(a). The clipped 3D geometry used to evaluate spatial gradients across the wound–healthy interface is shown in Figure 5(b). Fibroblast infiltration is slightly reduced, with *ρ/ρ*_0_ ≈ 1.51, due to the presence of pre-existing collagen within the wound region. This initial 40% collagen content provides additional mechanical support and partially counteracts the contractile forces generated during remodeling, thereby limiting inward dimpling of the wound region. By week 4, the cytokine concentration has decayed close to baseline, while the collagen density increases from *ϕ* = 0.40 to approximately *ϕ* = 0.63. The scaffold increases *J*_*p*_ from 0.41 to 0.65, corresponding to a reduction in permanent contracture from approximately 59% to 35%. Overall, the scaffold moderates excessive contracture and stabilizes the structural remodeling process.

#### 4.1.2. The role of the fiber re-alignment of the 3D bio-printed implant

The biochemical field variables are extracted at the center of the wound for the 40% collagen scaffold while varying the initial fiber orientation angle (0°, 15°, 30°, and 45°), as shown in Figure 6. As observed in panels (a)–(c), the fibroblast density, cytokine concentration, and collagen density exhibit nearly identical temporal evolution for all fiber orientation angles. This behavior arises because the governing equations for fibroblast migration, cytokine kinetics, and collagen deposition are formulated as functions of the scalar collagen density *ϕ*, rather than the fiber orientation tensor. Consequently, the initial alignment angle does not much influence cellular infiltration or biochemical signaling within the wound.

**Figure 6:**
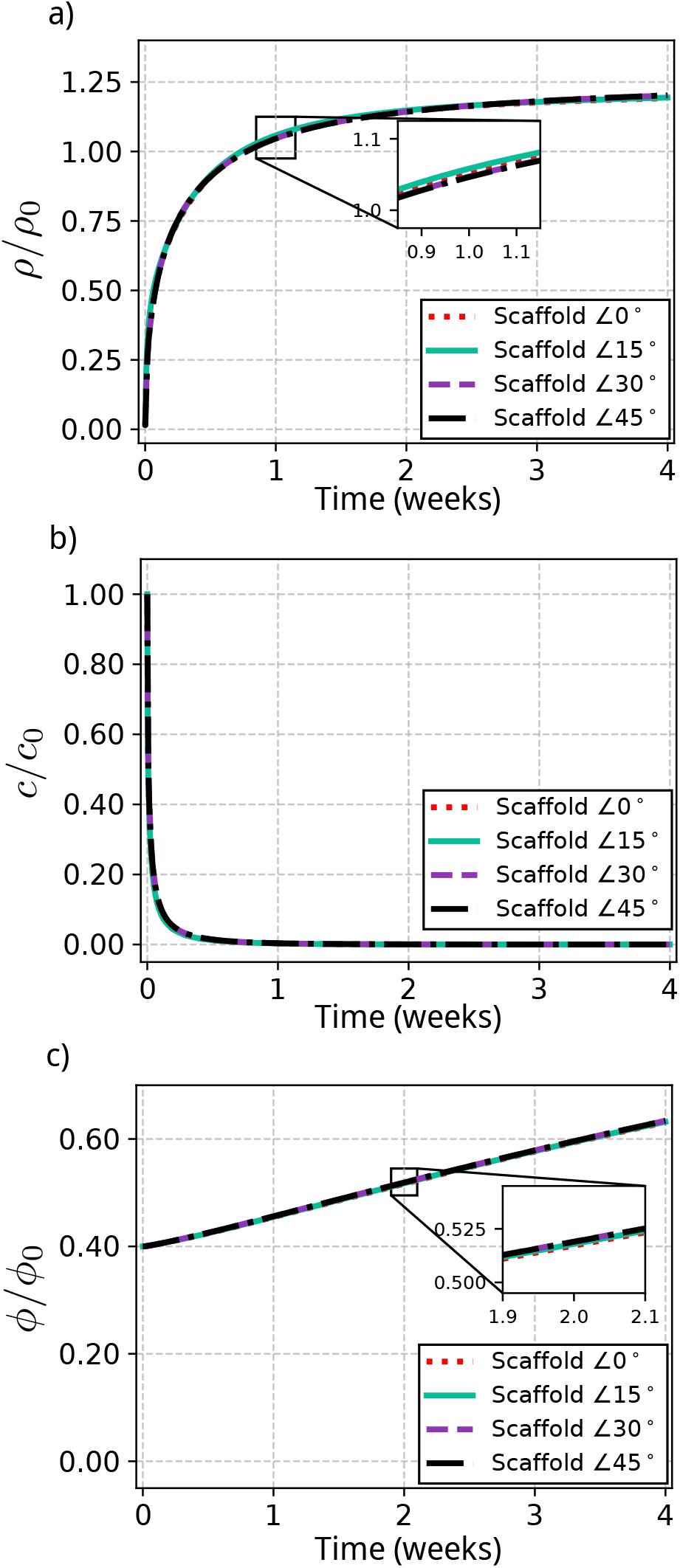
Evolution of (a) fibroblast infiltration, (b) cytokine concentration, and (c) collagen deposition at the wound-center showing over four weeks for endoscopic submucosal resection cases with a collagen fiber printing at different angles (0, 15, 30, 45).

Figure 7 shows the effect of scaffold fiber rotation on the mechanical response in the wound/scaffold region. The 0° case corresponds to the physiological diagonal fiber architecture, whereas 15°, 30°, and 45° represent progressively rotated scaffold fibers relative to this native configuration. Therefore, the angle variation should be interpreted as a local modification of scaffold anisotropy rather than a change in the global tissue architecture.

**Figure 7:**
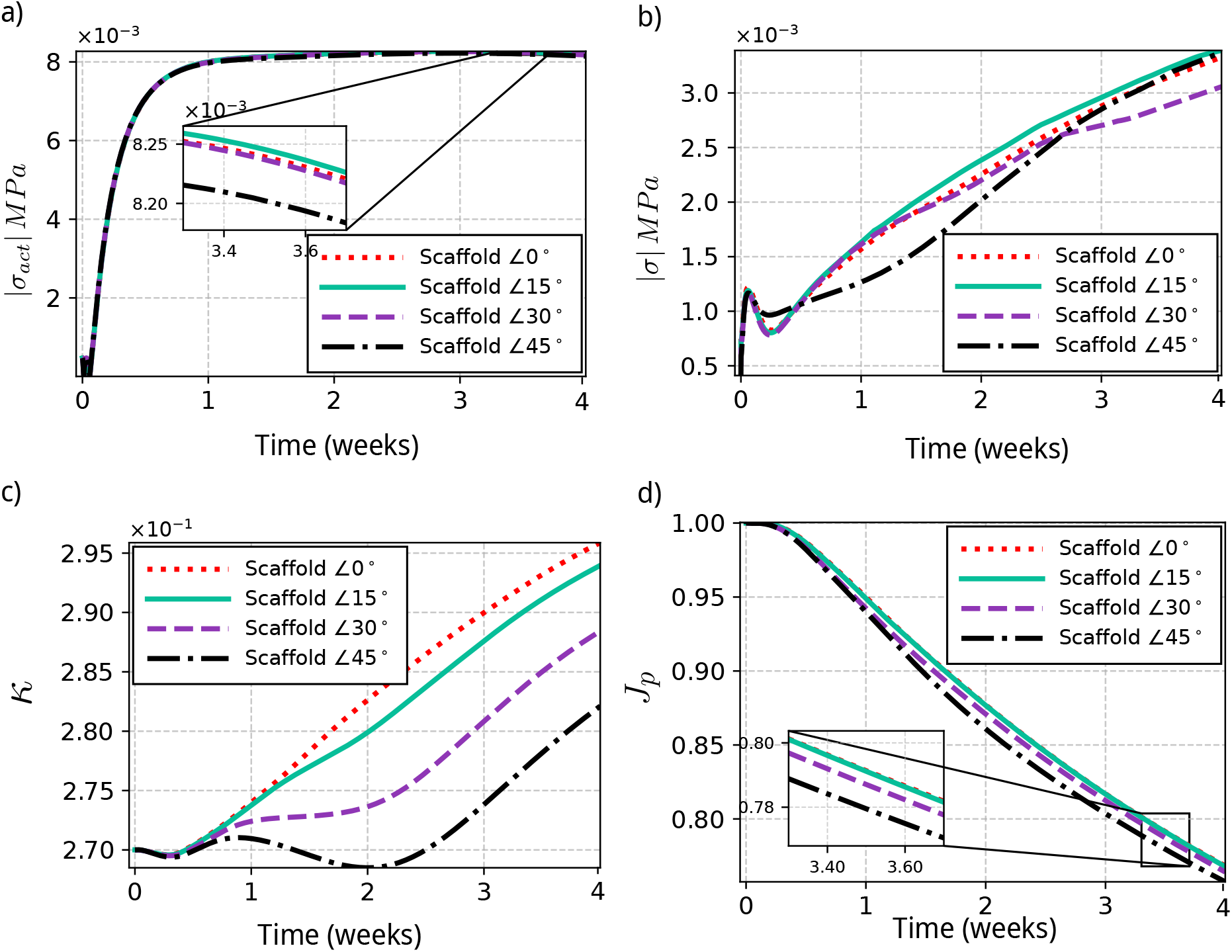
Evolution of the mechanical field variables, including (a) the active first Piola-Kirchhoff stress magnitude, (b) the total first Piola-Kirchhoff stress magnitude, (c) fiber dispersion (*κ* and the contracture *J*_*p*_ over a period of four weeks.

The active stress |*σ*_act_| as shown in Figure 7 (a) increases rapidly during the early healing phase and then approaches a nearly saturated response. This behavior is expected because |*σ*_act_| is mainly governed by fibroblast density, cytokine concentration, collagen content, and the evolving structure tensor. Since the biochemical fields are not directly prescribed by the scaffold angle, the active-stress curves remain relatively close. The slightly lower stress in the 45° case suggests that the contractile response is redistributed through the altered fiber architecture rather than amplified in magnitude.

The total passive stress |*σ*_total_| as shown in Figure 7 (b) exhibits a stronger sensitivity to scaffold orientation because it depends directly on the anisotropic fiber stretch invariants and the evolving collagen microstructure. Rotating the scaffold fibers changes their alignment with the local deformation field, leading to different levels of passive load transfer. The progressive increase in |*σ*_total_| reflects collagen deposition and remodeling, whereas the separation between angle cases indicates that scaffold orientation controls the mechanical resistance developed during healing.

The dispersion parameter *κ* as shown in Figure 7 (c) provides a mechanistic link between scaffold architecture and permanent remodeling. Lower values of *κ* indicate a more aligned and anisotropic fiber distribution, whereas higher values correspond to a more dispersed and isotropic network. The lower *κ* observed for larger scaffold rotations, particularly in the 45° case, suggests that these configurations promote a more directionally organized remodeling response. The dispersion of the fibers in the scaffold in No fill, 4%, 20% and 40% has 0.33, 0.314, 0.283 and 0.265 respectively. The use of scaffold-dependent dispersion parameters was motivated by the mechanobiological framework proposed by [23]. This means alignment with the fiber direction of the collagen submucosal layer **n**_*d*1_; of course, the speed of this alignment is conditioned by the initial fiber direction angle in the scaffold relative to the direction ***n***_*d*1_

The permanent deformation *J*_*p*_ shown in Figure 7 (d) decreases with increasing scaffold angle, indicating larger irreversible contracture in the wound/scaffold region. This response should not be interpreted as mechanically favorable; rather, it suggests a stronger scar-like contraction. The larger contracture in the 45° case does not contradict its lower |*σ*_act_|, because *J*_*p*_ is governed by accumulated plastic remodeling rather than active stress alone. The reduced *κ*, together with the altered passive anisotropic response, suggests that a more organized fiber network can transmit contraction more effectively into permanent deformation.

These results indicate that scaffold orientation regulates healing mechanics through the coupled evolution of active stress, passive anisotropic resistance, fiber dispersion, and plastic contracture. Increasing the scaffold angle promotes larger permanent contracture, which may be associated with an increased tendency toward scar-like tissue contraction. In contrast, the physiological 0° configuration provides a more balanced mechanical response with reduced irreversible contracture.

The full three-dimensional spatial distributions are examined to elucidate how fiber realignment governs tissue regeneration and structural remodeling. Figure 8 presents the spatial distributions of fibroblast density (*ρ*), cytokine concentration (*c*), collagen density (*ϕ*), and contracture (*J*_*p*_) for 40% collagen scaffold with an initial orientation 45° configuration at weeks 0 and 4. Owing to the presence of pre-existing collagen aligned at 45°, this results more permanent contracture compared to physiological alignment. The local contracture also creates deformation globally at week 4, as illustrated in Figure 8(a).

**Figure 8:**
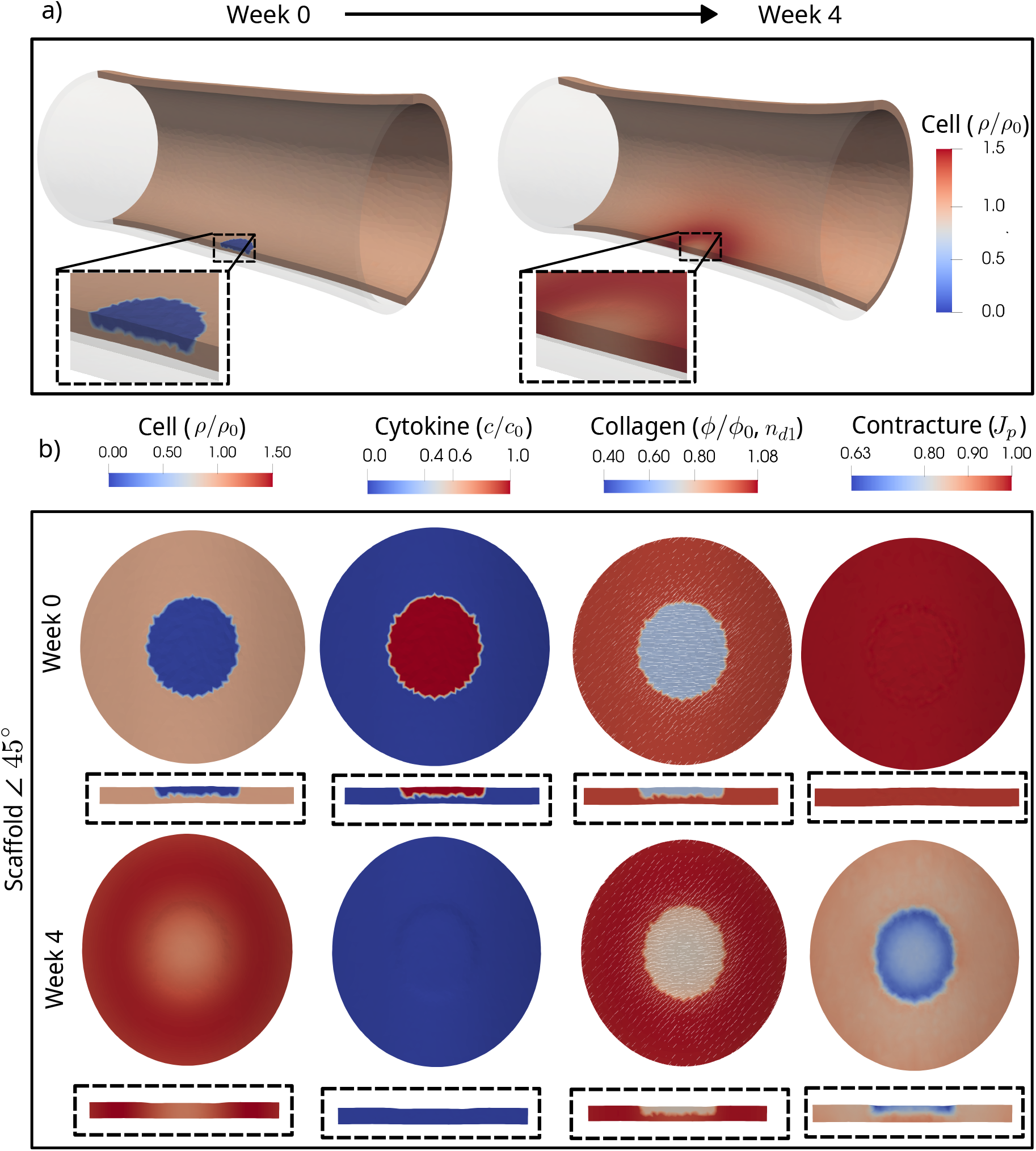
(a) Temporal solution illustrating changes in the wound region and cell production and infiltration around the region. (b) Extracted 3D subsection of the geometry showing the temporal evolution of fibroblast density, cytokine concentration, collagen density, and wound contracture for the 40 % scaffold with an initial fiber orientation of 20°, shown at weeks 0 and 4.

For the 40% collagen scaffold cases, changing the initial fiber orientation within the wound dressing produces only minor differences in the biochemical fields, indicating that fibroblast, cytokine, and collagen dynamics are not strongly affected by the prescribed scaffold angle, as illustrated in Figure 8(b). The collagen fibers, initially oriented at 45°, exhibit a gradual reorientation during healing, as illustrated in Figure 8(b). The main effect is instead mechanical, as reflected in the permanent deformation field *J*_*p*_. At week 4, the standard 40% scaffold gives *J*_*p*_ ≈ 0.65, whereas the 40%-45° scaffold gives *J*_*p*_ ≈ 0.63, indicating a modest increase in permanent contracture when the dressing fibers are initialized at 45° with respect to the homogenized diagonal fiber direction. This suggests that the initial scaffold orientation alters anisotropic load transfer and the subsequent redistribution of contractile stresses during remodeling.

Despite this slight increase in contracture relative to the standard 40% scaffold, the 40%-45° configuration still substantially reduces contracture compared with the no-fill case. The no-fill wound reaches *J*_*p*_ = 0.41 at week 4, corresponding to approximately 59% permanent contracture, whereas the 40%-45° scaffold reaches *J*_*p*_ ≈ 0.63, corresponding to approximately 37% contracture. Thus, the scaffold reduces permanent contracture by approximately 22% relative to the no-fill configuration but 40% scaffold has 24%. Overall, these results indicate that collagen scaffolding mechanically stabilizes the wound region and mitigates excessive scar-like contraction, while the initial fiber orientation further modulates the magnitude of local permanent remodeling.

### 4.2. Case study II: healing after intestinal end-to-end anastomosis

The end-to-end anastomosis case employs the same cylindrical geometry as previously described; however, the wound region is defined as a 10 mm wide circumferential annular band located at the mid-length of the domain. The initial and boundary conditions, as shown in Figure 9, were imposed in both the wound and the surrounding healthy regions. The simulation was carried out over a four-week healing period. The four fiber families were used for the anastomosis, as it is the resection of all fibers.

**Figure 9:**
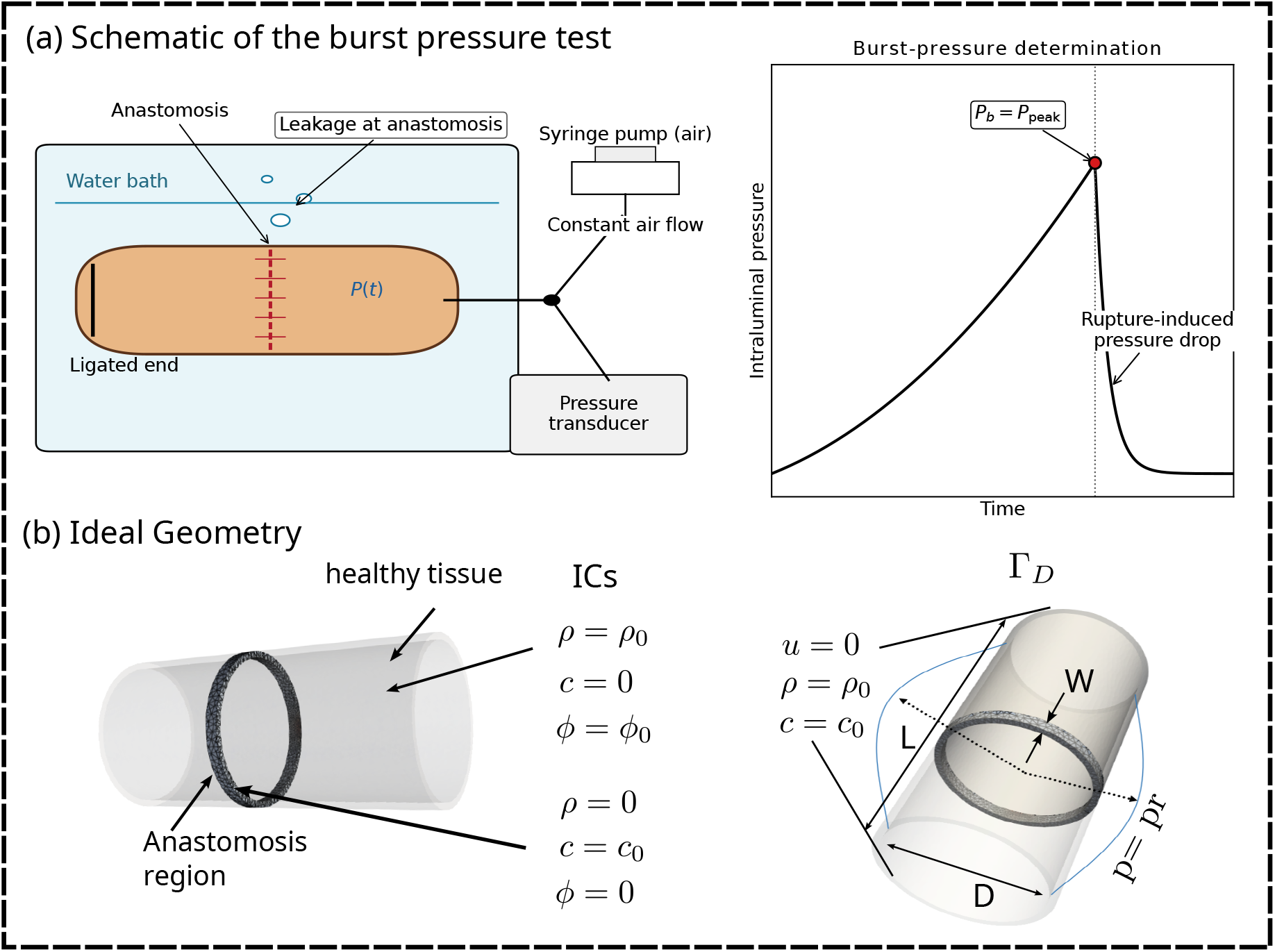
(a) Schematic representation of a commonly used experimental setup for burst pressure testing after intestinal anastomosis healing. Air is introduced into the healed intestinal specimen while it is submerged in water, allowing leakage or rupture to be visually identified by the appearance of air bubbles. In the corresponding numerical analysis, the burst event is identified by the sudden drop in pressure associated with leakage or rupture. (b) Idealized geometry with length (L), width (W), and diameter (D) considered for the anastomosis healing and burst pressure simulations.

The evolution of the biochemical field variables, including fibroblast density (*ρ*), cytokine concentration (*c*), collagen density (*ϕ*), and contracture (*J*_*p*_), at the center of the wound for different anastomosis damage widths is shown in Figure 10. The width of the anastomosis represents the extent of the tissue region affected by the anastomotic procedure and is varied to investigate its influence on the mechanobiological healing response.

**Figure 10:**
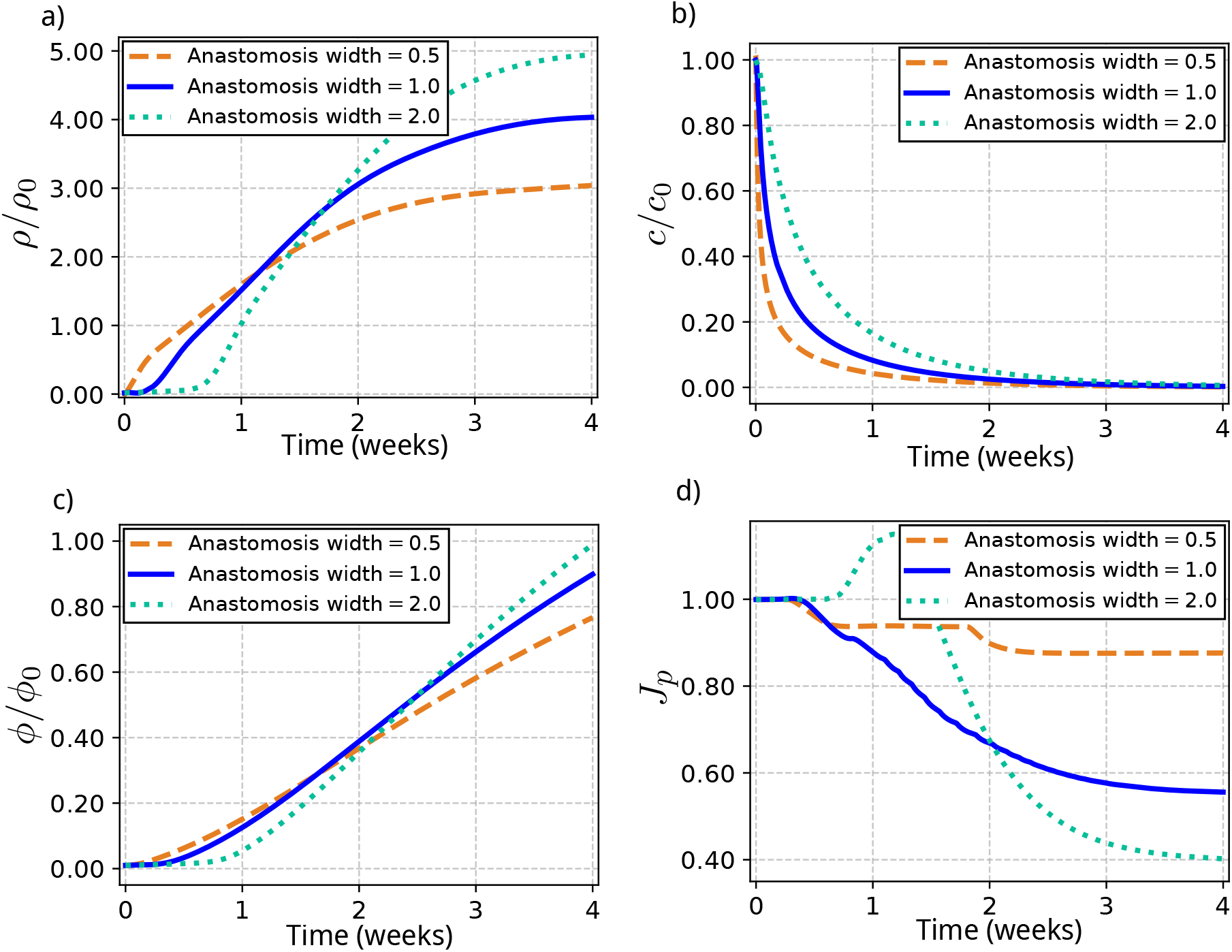
Temporal evolution at the wound-center location for the anastomosis cases with anastomosis region width 0.5, 1.0, 2.0, showing (a) fibroblast infiltration, (b) cytokine concentration, (c) collagen deposition over four weeks, and (d) Contracture over four weeks.

A smaller anastomosis width promotes faster diffusion and fibroblast proliferation, resulting in an earlier response and a lower fibroblast peak. In contrast, the exponential decay of cytokines is slower for wider approximately 1.0, as reduced cytokine-mediated degradation allows for eased fibroblast accumulation and prolonged cytokine response in wider regions, subsequently enhancing collagen deposition at later stages of healing. The final values of *J*_*p*_ for anastomosis widths of 0.5, 1.0, and 2.0 are approximately 0.87, 0.55, and 0.40, respectively. Thus, the reduction in *J*_*p*_ becomes more pronounced as the affected anastomotic width increases, indicating greater tissue contracture in wider regions. These results suggest that anastomotic techniques that minimize the width of the affected tissue region may reduce the extent of contracture during healing.

#### 4.2.1. Burst pressure (P_b_)

The burst pressure is defined as the minimum luminal pressure at which the mechanical driving stress exceeds the wound closure strength. Tissue failure is assumed to occur when the maximum principal stress induced by luminal inflation becomes greater than or equal to the wound closure strength generated by the total stress in the wound region. The burst pressure is therefore defined as:

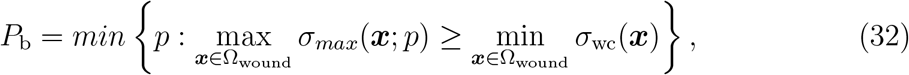

In Eq. (32) *σ*_*max*_ denotes the maximum principal Cauchy stress induced by luminal inflation during inflation test after healing, *σ*_wc_ represents the wound closure strength associated with total stress of healed tissue, and Ω_wound_ denotes the anastomosis region in the reference configuration.

The healing of the anastomosis case with width 1.0 mm was simulated, and monotonic inflation tests were performed from postoperative first hour to day 7. The wound closure strength increased as healing progressed, resulting in higher resistance to rupture. When the inflation-induced maximum principal stress reached the tissue strength threshold, as illustrated in Figure 11, up to postoperative day 6, failure occurred. By day 7, the wound closure strength exceeded the generated max stress by the applied inflation, and no burst was observed within the prescribed pressure range representing by “Burst Free” in Figure 11.

**Figure 11:**
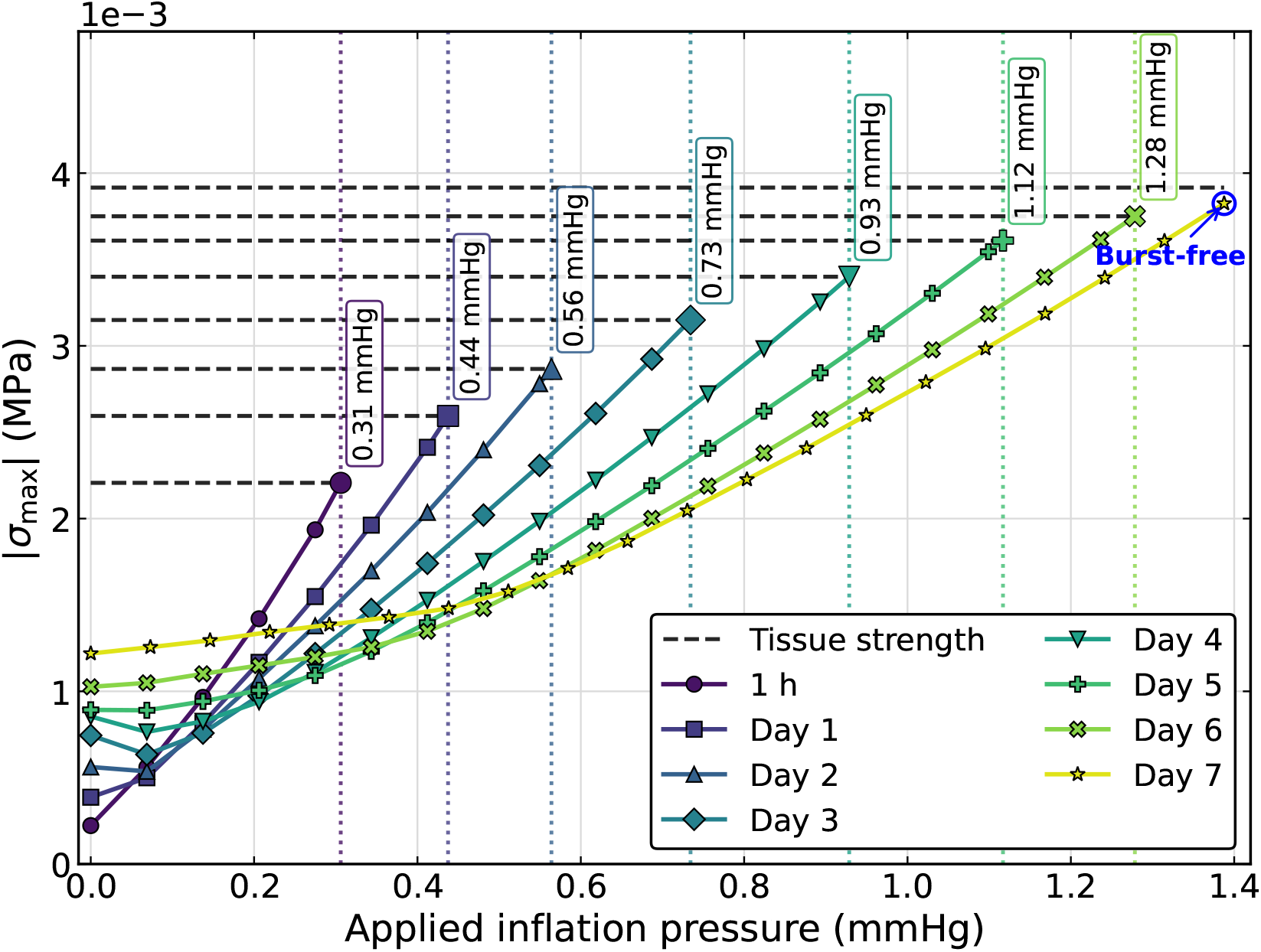
Applied Inflation pressure (mmHg) versus maximum principal stress magnitude (MPa). As healing progresses, tissue strength increases due to collagen and remodeling, resulting in higher burst pressures. By day 7, the tissue remains burst-free within the tested pressure range (blue circle).

Figure 12(a) shows the spatial distribution of collagen density and active stress in the anastomotic region at postoperative days 3 and 7. The zoomed cross-sectional view highlights localized wall thinning at day 3, associated with reduced collagen density and consequently lower structural stiffness. By day 7, collagen deposition increased within the wound region, leading to partial restoration of wall thickness and improved mechanical integrity.

**Figure 12:**
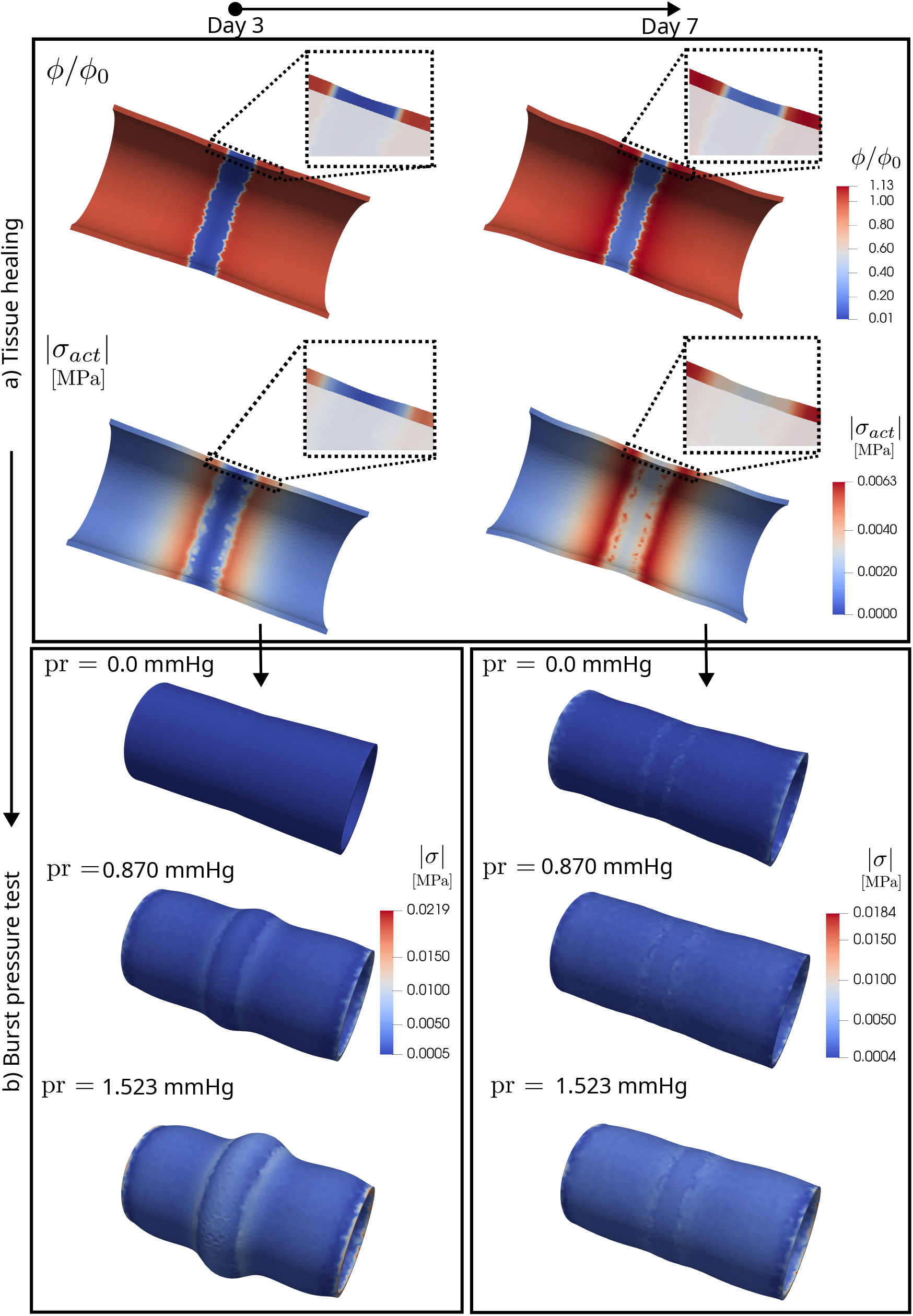
(a) Distribution of collagen density and active stress during tissue healing at postoperative days 3 and 7, including a zoomed view of the anastomotic region illustrating wall thinning. The color bar indicates the magnitude of the respective fields. The same minimum–maximum range is used for both time points to ensure that the values lie within the same scale. Colors represent collagen density ranging from low (dark blue) to high (light blue). (b) Results of the burst pressure test under the same inflation protocol for the anastomosis case at postoperative days 3 and 7.

The active stress distribution exhibited reduced magnitude near the anastomotic edge at day 3. At day 7, the stress became more uniformly distributed with higher concentration along the anastomotic interface, indicating enhanced load-bearing capacity of the healed tissue. After healed tissue at days 3 and 7, burst pressure tests were performed by applying monotonic inflation while maintaining the corresponding collagen density field, as shown in Figure 12(b). The burst pressure was defined as the inflation pressure at which the maximum principal stress exceeded the wound closure strength threshold as shown in Eq. (32). At day 3, the reduced collagen density resulted in lower stiffness, leading to greater localized deformation and earlier attainment of the burst criterion. In contrast, at day 7, increased collagen deposition enhanced tissue stiffness and resistance to inflation, thereby improving structural stability at the anastomotic region. The day 7 healing case did not reach the burst threshold under the applied loading conditions.

The temporal evolution of burst pressure from postoperative first hour to day 7 is presented in Appendix G. The Figure G.19, showing progressive recovery of tissue strength, with the healed tissue exceeding the burst pressure requirement by day 7. There is a strong correlation of burst pressure and collagen deposition. We can therefore conclude that, as collagen is progressively deposited over time, it contributes to strengthening the tissue in the anastomosis region, ultimately improving its resistance to the intraluminal pressure applied during burst testing.

## 5. Conclusion

A mechanobiological formulation was adapted to model wound healing in the GI tract, accounting for multiple fiber families of reinforcement and layered architecture. The framework couples reaction–diffusion dynamics with nonlinear finite strain mechanics, incorporating isotropic, anisotropic, and volumetric strain-energy contributions. A multiplicative decomposition separates elastic and plastic deformation, while active stress captures contractility-induced deformations. Collagen production, remodeling, fiber dispersion, and plastic stretch modulate tissue stiffness and anisotropic response. The governing equations were discretized using finite elements in space and backward Euler in time and solved through a staggered strategy in `FEniCS`.

In the ESD configuration, higher initial collagen content limited fibroblast infiltration and reduced plastic contracture compared with lower-density scaffolds. Variations in collagen fiber orientation minimally affected biochemical evolution and total collagen production but significantly influenced the structural response: greater deviation from physiological fiber directions increased contracture through anisotropic stiffness and remodeling effects. The anastomosis configuration exhibited elevated fibroblast density and collagen accumulation, highlighting the influence of wound topology on biochemical–mechanical coupling and stress redistribution. Predicted burst pressure strongly correlated with collagen density, linking collagen deposition to mechanical recovery. Plastic stretch further indicated that a portion of wound contraction was irreversible, reflecting permanent remodeling during late-stage healing.

The framework enables quantitative assessment of healing under variations in ESD scaffold density and fiber orientation and extends to anastomotic healing and burst pressure prediction. These findings provide a computational basis for investigating how matrix composition, fiber architecture, and wound configuration influence GI healing and mechanical recovery.

## 6. Limitations and perspectives

The present formulation assumes a homogenized fiber distribution across the intestinal wall thickness, thereby neglecting the explicit layer-wise organization of fibers. A thickness-resolved representation could enable more accurate characterization of layer-specific contracture and facilitate improved assessment of healthy and pathological conditions such as stenosis [16], where pathologically enhanced circumferential contraction may contribute to luminal narrowing in the anastomotic region. Future work could incorporate collagen-fiber remodeling through selective deposition and removal [44], linking collagen turnover with evolving fiber orientation, dispersion, and tissue anisotropy during healing.

The predictive capability of the framework is constrained by scarce quantitative intestinal healing data, particularly fibroblast and collagen density with cytokine concentration, and mechanobiological parameters. Experimental characterization would improve the identification of intestine-specific parameters and support subject-specific healing predictions. Burst pressure characterizes anastomotic mechanical integrity, but rupture is inferred from a threshold based on evolving wound closure strength rather than an explicit failure mechanism. Incorporating a continuum damage or rupture internal variable could enable more mechanistic predictions of burst pressure and progressive loss of load-carrying capacity.

## Acknowledgments

The authors acknowledge support from the Italian National Group for Mathematical Physics (GNFM–INdAM) and the European Union’s Horizon Europe research and innovation programme under grant agreement No. 101170592 (MiGEM). The authors also acknowledge CINECA for providing high-performance computing resources through the ISCRA Class C project GI-HEAL (HP10CA591G).

## Appendix A Mechanical parameter estimation

The accurate identification of mechanical parameters is essential for capturing the passive behavior of the healing colon model. To calibrate the constitutive parameters, four fiber families were considered together with a single dispersion parameter. Several experimental studies have been conducted on intestinal tissues under various loading conditions, including uniaxial [29], biaxial [45], cylinder occlusion [28], and inflation–extension tests [46], to characterize their mechanical response.

In the present study, no new experimental measurements were performed. Instead, experimental data points were extracted from the literature using the online tool Plot-Digitizer. The parameter optimization procedure was implemented in Python using the lmfit library. Unlike the assumption in Patel et al. [46], where the circumferential fiber family was neglected due to its smaller contribution, all four fiber families were retained in this work. It was further assumed that the healthy dispersion parameter, *κ*, remained identical across all fiber families. The angle range was initially set from 30° to 50° for parameter estimation [47].

The material parameters were identified by fitting the experimental pressure–outer diameter and axial force–outer diameter data using the following integral relations for pressure and axial force:

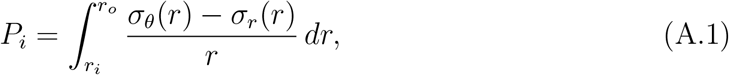

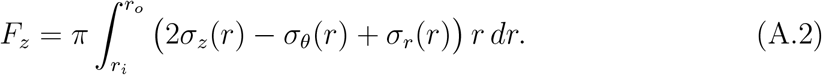

**Figure A.13.**
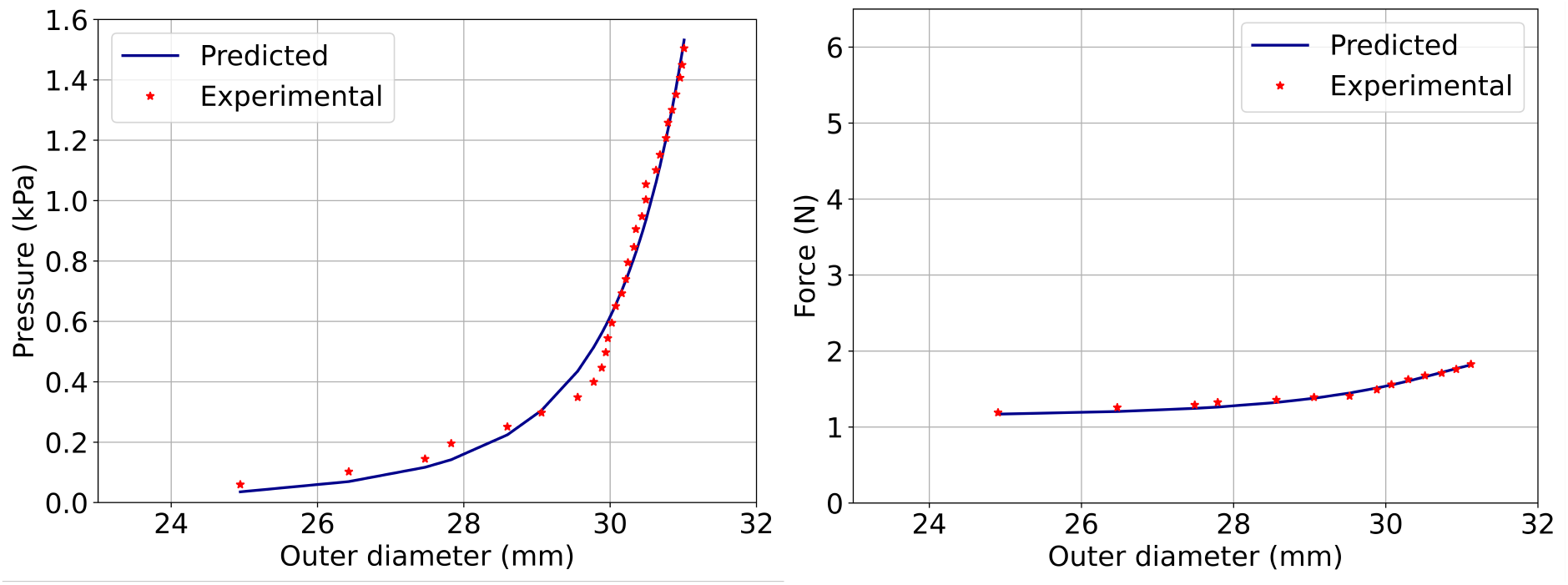
Estimation of mechanical parameters by fitting the pressure–outer diameter and force–outer diameter curves to the experimental inflation–extension data.

**Table A.1:** Estimated mechanical parameters for the passive behavior of the healing colon model.

| Parameter | $k_0$ [kPa] | $k_v$ [MPa] | $k_1^l$ [kPa] | $k_1^c$ [kPa] | $k_1^{d1}$ [kPa] |
| --- | --- | --- | --- | --- | --- |
| Value | 2.10 | 0.32 | 3.43 | 0.78 | 2.50 |
| Parameter | $k_2^l$ [-] | $k_2^c$ [-] | $k_2^{d1}$ [-] | $\kappa$ [-] | $\theta$ [$^\circ$] |
| Value | 1.09 | 0.02 | 0.31 | 0.23 | 44.91 |

## Appendix B. Fiber remodeling validation

### Appendix B.1. Remodeling of soft biological tissue strip under tension

The benchmark test for the validation of soft tissue remodeling is performed on the strip under tension [38]. In this test, a rectangular domain of length *L* = 4 mm and height *H* = 1 mm is considered. The left side of the domain is fixed, while on the right side, a quasi-static constant displacement Δ*u* = 1 mm on the *x*-axis. The material has an embedded family of fibers described by the unit vector ***n***_*f*_ having an angle *α* = 45° with respect to the *y*-axis. The constitutive material behaviour is described by a strain energy density of the following form Ψ = Ψ^iso^ + Ψ^aniso^ of Neo-Hookean type:

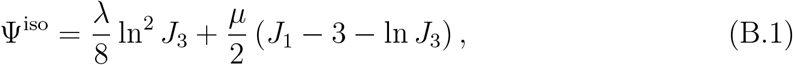

and

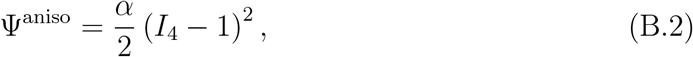

where *J*_1_ = tr (***C***), *J*_3_ = det(***C***), *I*_4_ = tr (***CA***) and the structural tensor ***A*** is given by ***A*** = ***n***_*f*_ ⊗ ***A***_*f*_. The remodeling of the fibers in the direction of the principal stretch 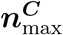 of ***C*** follows the equation:

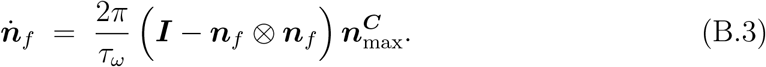

**Figure B.14:**
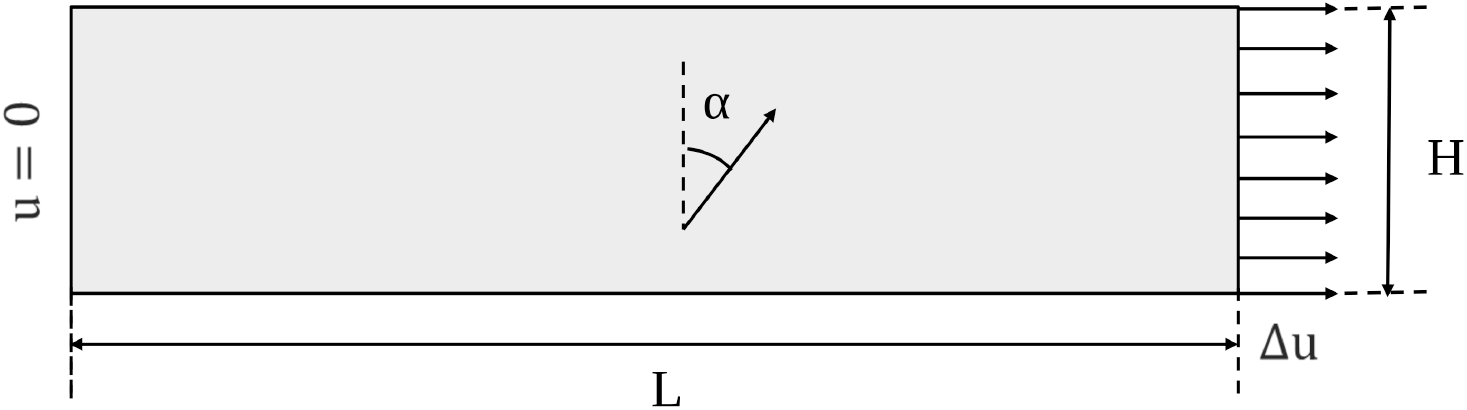
Schematic of the rectangular strip specimen with length 4 mm and height 1 mm. Collagen fibers are initially oriented at an angle *α* = 45° with respect to the vertical axis. A horizontal displacement Δ*u* is prescribed at one end of the strip, while the opposite end is fully fixed.

**Figure B.15:**
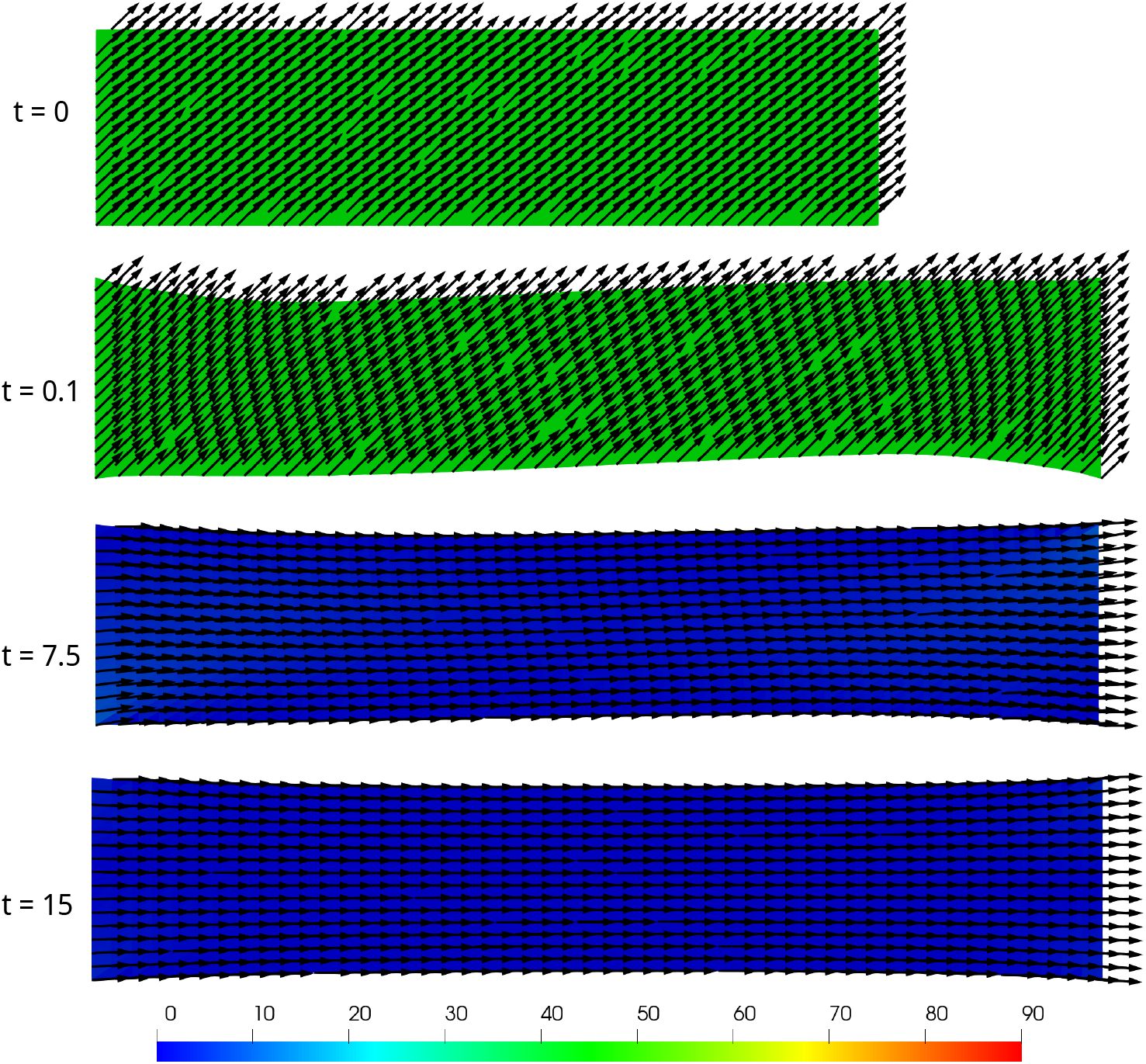
The fiber ***n***_*f*_, initially oriented at 45°, undergoes stretch-driven reorientation toward the maximum principal stretch direction 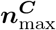. The orientation progressively decreases from 45° (green) to 0°, indicating full alignment (blue).

### Appendix B.2. Cylindrical tube subjected to internal radial displacement

A sinusoidal radial displacement was prescribed in the inner wall, with both ends fixed. The resulting principal stretch drove the fibers to reorient predominantly in the circumferential direction at the center of the tube.

**Figure B.16:**
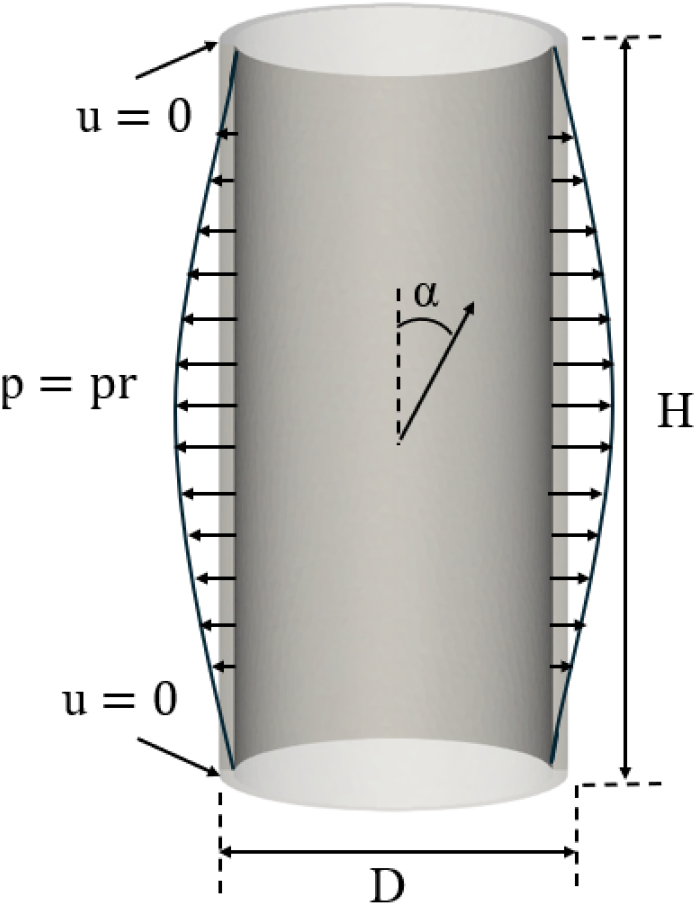
Schematic diagram for radial displacement load.

**Figure B.17:**
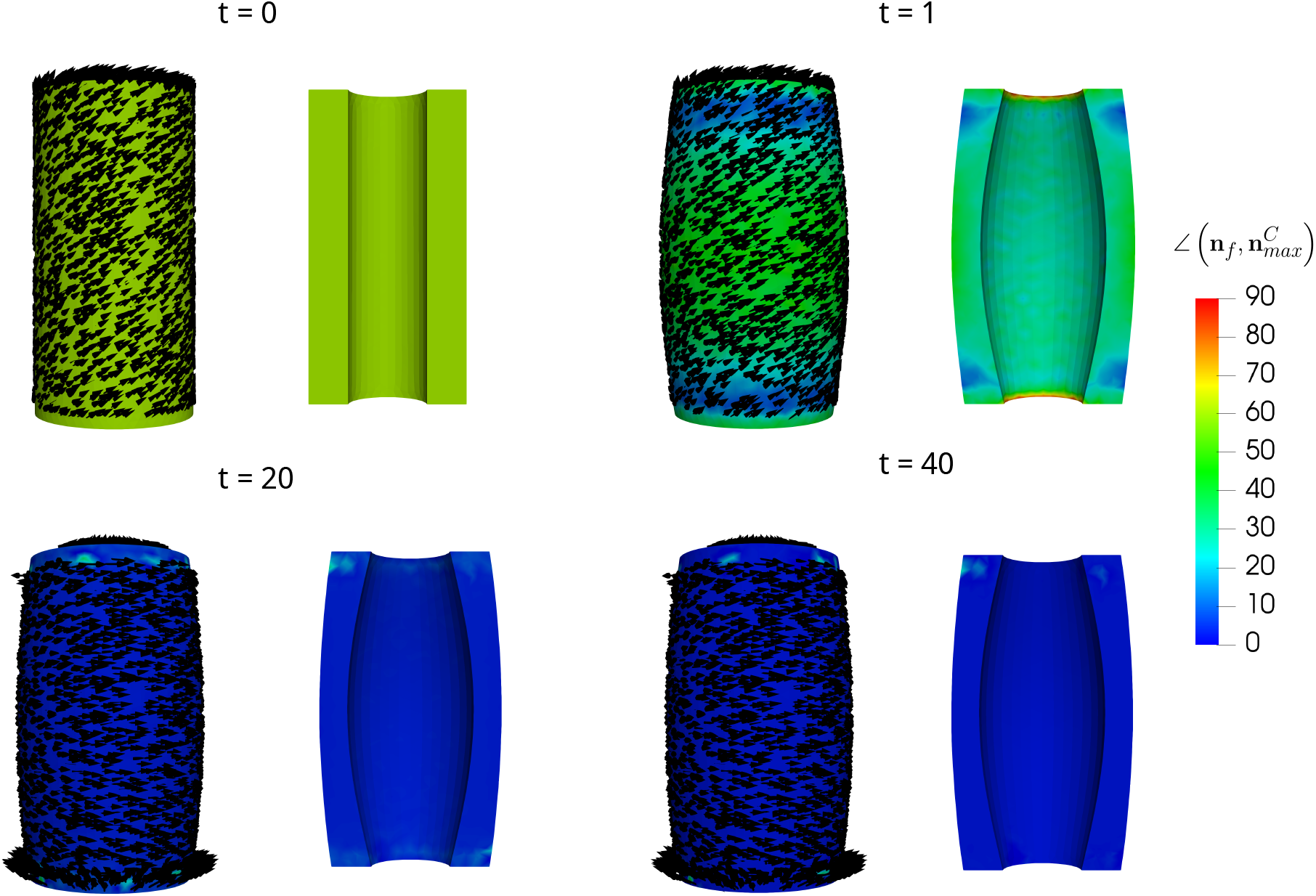
The fiber ***n***_*f*_, initially oriented at 60°, undergoes inflation-driven reorientation toward the maximum principal stretch direction 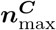. The orientation progressively decreases from 60° to

## Appendix C. Time integration

### Appendix C.1. Mechanical Weak form residual

The residual of the weak mechanical problem from equations (20), together with the Neumann and Dirichlet boundary conditions in Eqs. (22) and (21), respectively, is defined as:

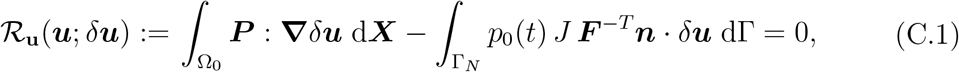

for all *δ****u*** ∈ V_0_, where V_0_ incorporates the homogeneous Dirichlet conditions on Γ_*D*_.

### Appendix C.2. Biochemical weak form residual

The implicit backward Euler time integration scheme is applied in a weak form in equations (26) and (27), and the corresponding residuals are defined as:

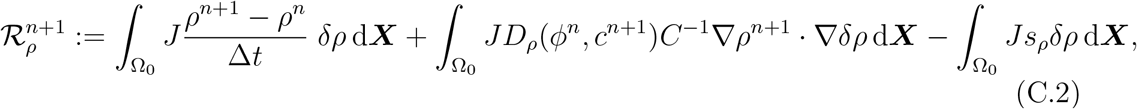

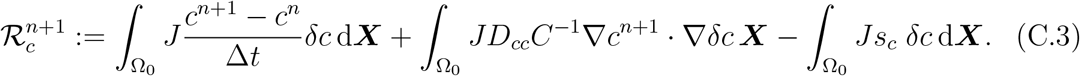

In compact notation, the discrete problem to be solved in each timestep *t*^*n*+1^ is: Find (*ρ*^*n*+1^, *c*^*n*+1^) such that:

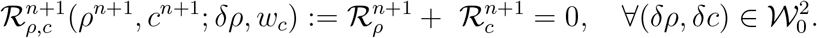

### Appendix C.3. Micro-structural field weak form residual

The weak form equations (28), (29), (30), and (31) are discretized using the implicit backward Euler method and written down in residual form as:

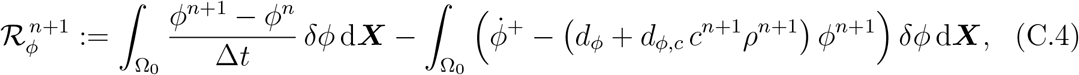

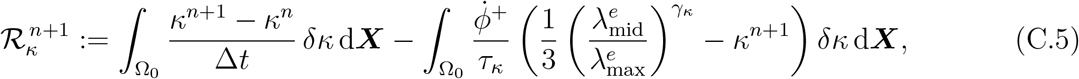

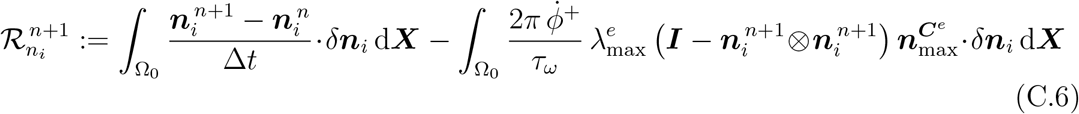

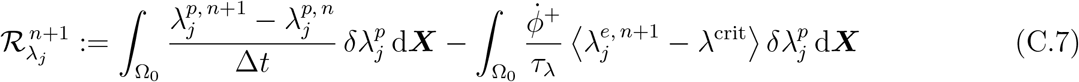

The discrete microstructural residuals at *t*^*n*+1^ are assembled into a single global nonlinear residual:

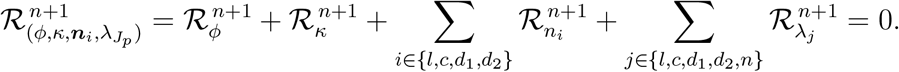

## Appendix D. Algorithm

### Algorithm 1

Algorithm for the mechano-biological healing of the GI tract.

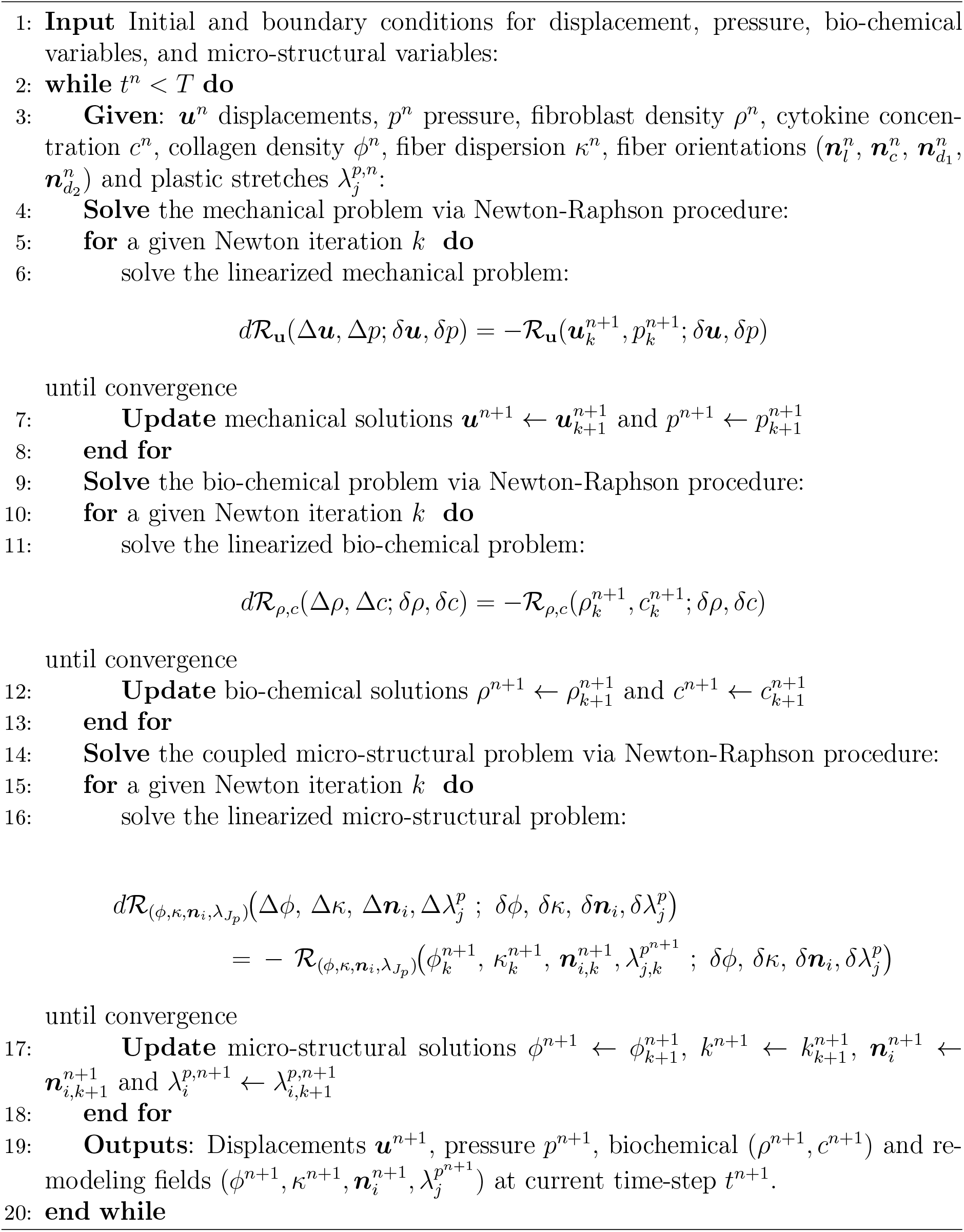

## Appendix E. Mechanobiological Parameters

The mechanobiological parameters used in the healing model are as follows:

**Table E.2:** Cells and cytokines population dynamics parameters.

| Parameter | Value | Interpretation | Ref. |
| --- | --- | --- | --- |
| $p_\rho [\text{h}^{-1}]$ | $9.0068 \times 10^{-4}$ | Fibroblast proliferation rate. | [23] |
| $p_{\rho,c} [\text{h}^{-1}]$ | 0.015314 | Cytokine-mediated fibroblast proliferation | [5] |
| $K_{\rho,c} [-]$ | $1.0 \times 10^{-5}$ | Saturation constant for cytokine-induced proliferation. | [5] |
| $K_{\rho\rho} [-]$ | 1000.0 | Fibroblast division saturation | [48] |
| $d_\rho [\text{h}^{-1}]$ | $8.1061 \times 10^{-4}$ | Effective fibroblast loss rate. | [48] |
| $d_{\rho,c} [\text{mm}^2\text{hr}^{-1}]$ | $6.12 \times 10^{-3}$ | Cytokine increase fibroblasts diffusivity | [49] |
| $d_{\rho,0} [\text{mm}^2\text{hr}^{-1}]$ | $6.12 \times 10^{-5}$ | Baseline fibroblast diffusivity | [31] |
| $d_{\rho,\phi} [\text{mm}^2\text{hr}^{-1}]$ | $1.58 \times 10^{-3}$ | Cross-diffusion between fibroblasts and cytokines. | [24] |
| $p_{\rho,J_e} [\text{h}^{-1}]$ | $4.503 \times 10^{-4}$ | Mechanically induced fibroblast activation rate. | [23] |
| $\rho_0 [\text{cells}/\text{mm}^3]$ | 55051.26 | Maximum fibroblast density in healthy tissue. | [24] |
| $c_0 [\text{g}/\text{mm}^3]$ | $1.0 \times 10^{-4}$ | Initial cytokine concentration. | [5] |
| $p_{c,\rho} [\text{h}^{-1}]$ | $1.63 \times 10^{-18}$ | Cytokine production induced by fibroblasts. | [5] |
| $p_{c,J_e} [\text{h}^{-1}]$ | $5.45 \times 10^{-18}$ | Cytokine production induced by mechanical activation. | [5] |
| $K_{c,c} [\text{mol}/\text{mm}^3]$ | 1.0 | Cytokine saturation | [5] |
| $d_c [\text{h}^{-1}]$ | 0.005 | Cytokine degradation rate. | [50] |
| $D_c [\text{mm}^2\text{h}^{-1}]$ | 0.01208 | Diffusion coefficient of cytokines. | [51] |

**Table E.3:** Tissue local remodeling and traction parameters.

| Parameter | Value | Interpretation | Ref. |
| --- | --- | --- | --- |
| $p_\phi [\text{h}^{-1}]$ | $1.4 \times 10^{-8}$ | Baseline collagen production rate. | [52] |
| $p_{\phi c} [\text{h}^{-1}]$ | $7.0 \times 10^{-8}$ | Cytokine-induced collagen production rate. | [51] |
| $p_{\phi, J_e} [\text{h}^{-1}]$ | $1.4 \times 10^{-8}$ | Mechanically induced collagen production rate. | [5] |
| $K_{\phi c} [-]$ | $1.0 \times 10^{-4}$ | Cytokine saturation constant for collagen synthesis. | [5] |
| $d_\phi [\text{h}^{-1}]$ | $3.7413 \times 10^{-4}$ | Collagen degradation | [52] |
| $d_{\phi c} [\text{h}^{-1}]$ | $8.81 \times 10^{-6}$ | Cytokine-induced collagen degradation | [52] |
| $K_{\phi, \rho} [-]$ | 1.06 | Collagen saturation constant controlling remodeling dynamics. | [5] |
| $\gamma_{je} [-]$ | 5.0 | Shape of mechanical curve | [5] |
| $\theta_e [-]$ | 2.0 | Mechanical activation threshold. | [53] |
| $\tau_w [\text{h}]$ | 4.8543 | Time constant for fiber reorientation. | [5] |
| $\tau_k [\text{h}]$ | 0.4854 | Time constant for fiber dispersion evolution. | [5] |
| $\gamma_k [-]$ | 5.0 | Shape of dispersion rate curve | [5] |
| $\tau_{p(l, c, d_1, d_2, n)} [\text{h}]$ | 0.05 | Plastic relaxation time (circumferential, longitudinal, diagonals, and normal direction). | [5] |
| $t_p [\text{MPa}]$ | $3.503 \times 10^{-8}$ | Baseline active stress generated by fibroblasts. | [5] |
| $t_{pc} [\text{MPa}]$ | $1.051 \times 10^{-7}$ | Up-regulation of contraction by the presence of cytokines | [5] |
| $K_{tc} [-]$ | $1.0 \times 10^{-5}$ | Cytokine saturation constant for active stress generation. | [5] |
| $K_t [-]$ | 0.4 | Saturation constant controlling active stress response. | [5] |

## Appendix F. Initial fiber configuration

The schematics of the wound geometry for the no-fill case and scaffold-based wound dressings with aligned and with fibers oriented at 45° relative to the healthy tissue fibers are considered in the ESD simulations, as shown in Figure. F.18.

**Figure F.18:**
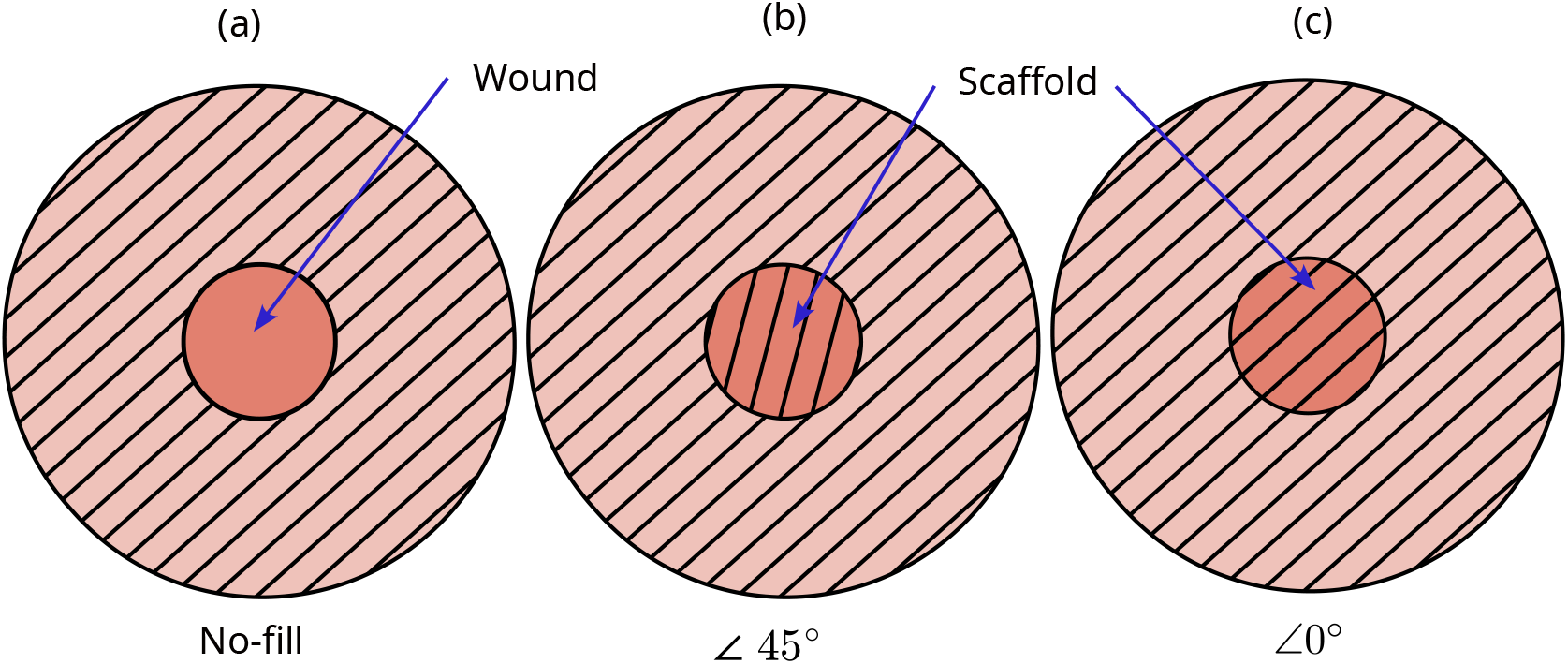
Initial fiber distribution in the wound region for the bioprinted implants: (a) no fill; (b) scaffold fibers oriented at ∠45° relative to the collagen fibers of the surrounding healthy tissue; (c) scaffold fibers aligned (∠0°) with the collagen fibers of the surrounding healthy tissue.

## Appendix G. Burst pressure evaluation

Burst pressure was evaluated using a failure criterion in which rupture occurs when the maximum principal stress induced by luminal inflation equals or exceeds the tissue strength. The role of collagen production is important to improve burst pressure.

Figure G.19 shows a strong positive association between collagen density and burst pressure (*r* = 0.960, *n* = 7). Day 7 was excluded because no burst occurred at the maximum applied pressure, giving only *P*_burst_ *>* 1.38 mmHg. This association suggests that collagen accumulation contributes to the recovery of anastomotic strength.

**Figure G.19:**
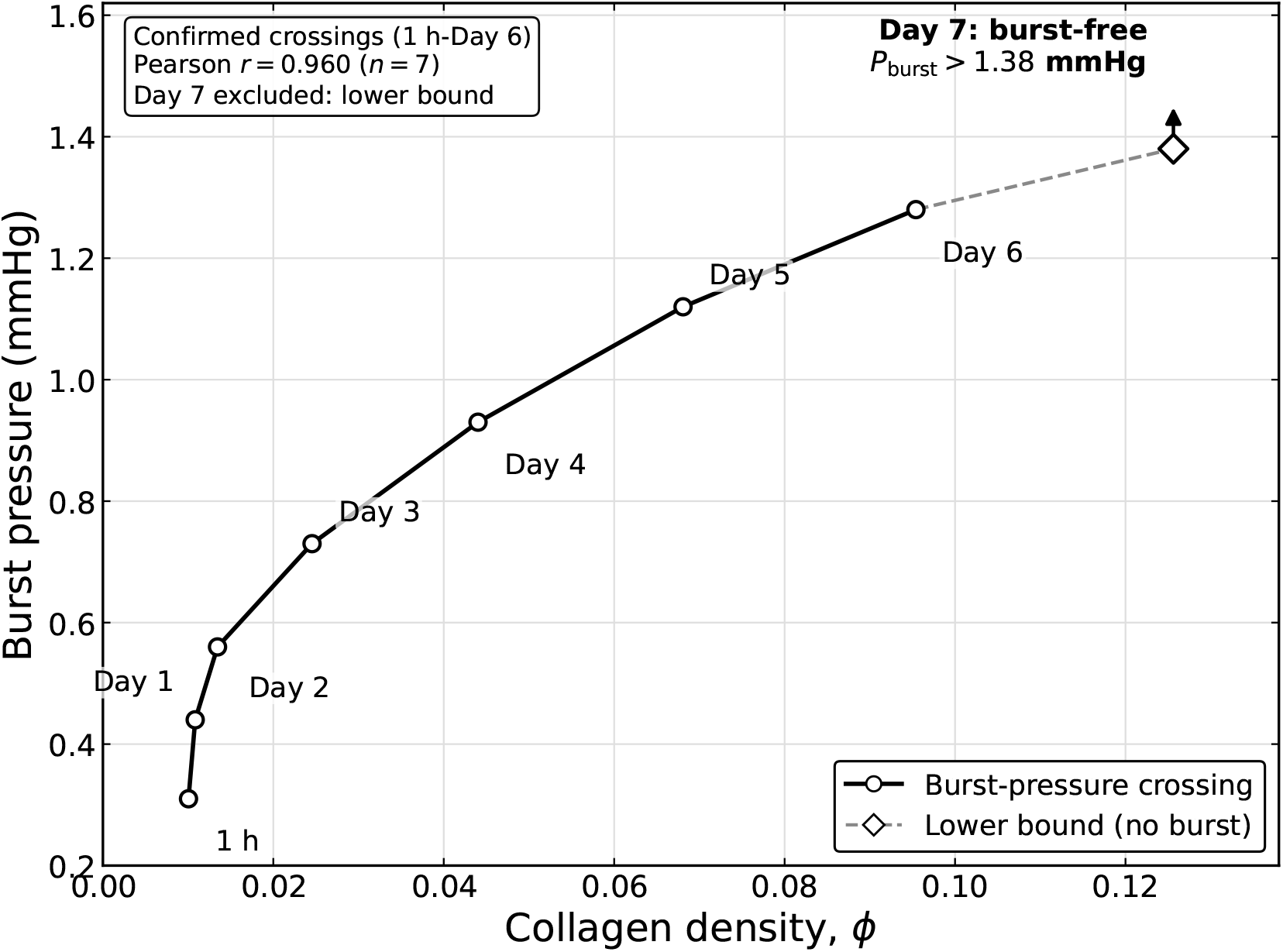
Relationship between collagen density and burst pressure. Day 7 represents a lower bound because no burst occurred at the maximum applied pressure.

## Appendix H. Remodeling of intestinal fibers

Figure H.20 shows fiber remodeling for the 40% collagen scaffold with initial angles of 0° (a) and 45° (b) at weeks 0, 2, and 4. The glyphs represent the local fiber directions, while the colors indicate their angles relative to the physiologically healthy tissue fiber direction. In both cases, the fibers reorient toward the evolving maximum principal stretch direction. Continued remodeling and contracture alter this direction, particularly near the wound boundaries, resulting in spatially nonuniform fiber orientations. The increase in the angle of the scaffold results in more contracture and delay in remodeling.

**Figure H.20:**
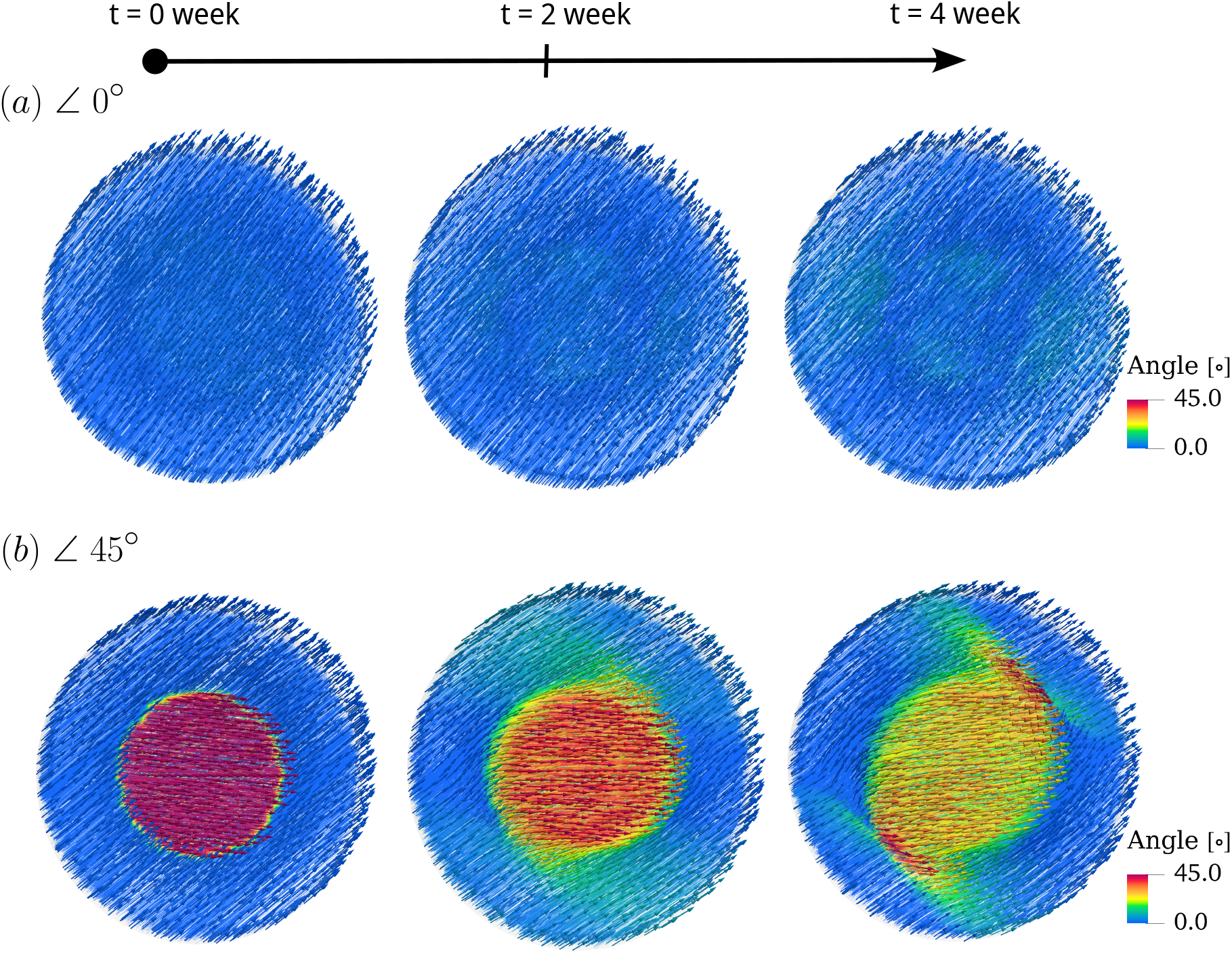
Evolution of diagonal fiber orientation from an initial alignment with 0° and with angle 45° toward the physiological reference direction.

## Notes

### Competing Interest Statement

The authors have declared no competing interest.

